# Kinematic performance declines as group size increases during escape responses in a schooling coral reef fish

**DOI:** 10.1101/2023.09.15.557889

**Authors:** Monica D. Bacchus, Paolo Domenici, Shaun S. Killen, Mark I. McCormick, Lauren E. Nadler

**Author notes:** **Correspondence:** Monica D. Bacchus, Lauren E. Nadler.

## Abstract

Escaping predation is essential for species survival, but prey must effectively match their response to the perceived threat imposed by a predator. For social animals, one mechanism to reduce risk of predation is living in larger group sizes, which dilutes each individual’s risk of capture. When a predator attacks, individuals from a range of taxa (e.g., fishes, sharks, amphibians) perform an escape response, to evade the attack. Here, using the schooling coral reef damselfish *Chromis viridis*, we assess if there is an optimal group size that maximizes both individual escape response performance as well as group cohesion and coordination following a simulated predator attack, comparing schools composed of four, eight, and sixteen fish. We found that fish in various group sizes exhibited no difference in their reaction timing to a simulated predator attack (i.e., escape latency), but larger groups exhibited slower kinematics (i.e., lower average turning rate and shorter distance covered during the escape response), potentially because larger groups perceived the predator attack as less risky due to safety in numbers. Both school cohesion and coordination (as measured through alignment and nearest neighbor distance, respectively) declined in the 100ms after the predator’s attack. While there was no impact of group size on alignment, larger group sizes exhibited closer nearest neighbor distances at all times. This study highlights that larger group sizes may allow individuals to save energy on costly behavioral responses to avoid predators, potentially through a greater threshold of the threat necessary to trigger a rapid escape response.

## 1 Introduction

Numerous fish species use social behavior as a strategy to reduce the risk of predation and share the costs associated with predator vigilance (Krause and Ruxton, 2002;Ward and Webster, 2016). An individual’s vulnerability to predation depends on factors such as the predator’s ability to detect the group, the predator’s attack rate, and the likelihood of the individual escaping the attack (Roberts, 1996). Compared to individuals in smaller shoals, individuals in larger shoals benefit from dilution of individual risk of predation but also face a higher risk of being spotted by predators due to increased visibility (Krause and Godin, 1995;Killen et al., 2017). When a predator is detected, shoals collectively adapt their response based on available sensory and social information (Brown et al., 2006;Marras et al., 2012;Rieucau et al., 2014a), with socially transmitted information communicated more effectively when the group exhibits a higher level of organization and greater connectedness between neighbors (Rosenthal et al., 2015).

However, larger group sizes also come with a tradeoff between the defensive benefits they offer and the increased competition for limited resources like food, habitat, and breeding partners (Hoare et al., 2004;Gil et al., 2017). Social fishes may modify their coordination (e.g., polarized alignment) and cohesion (e.g., distance to the nearest neighbor) based on the context. For example, hungrier fish have been shown to become less cohesive, possibly to prioritize individual foraging (Morgan, 1988;Krause, 1993). Conversely, shoals enhance cohesion and coordination when facing threats (Herbert-Read et al., 2017). These tradeoffs lead to an optimal group size that balances the costs of group living against its potential benefits (Brown, 1982;Pulliam and Caraco, 1984;Killen et al., 2017). In resource-scarce habitats, smaller groups may be favored over larger groups to minimize competition (Krause and Ruxton, 2002). Conversely, larger groups are likely preferred over smaller groups when predation risk is high (Hager and Helfman, 1991). Ward & Webster (2019) found that mid-sized groups (composed of 12-20 fish) of the three-spined sticklebacks (*Gasterosteus aculeatus*) balanced decisions related to foraging and risk better than either the smallest (with as few as two fish) or largest group sizes (composed of up to 29 fish) tested. Yet, we know little about the effect of group size on the response to an attacking predator in the crucial first few milliseconds after a predator attacks.

During predator attacks, fishes employ escape responses to swiftly move away from danger, increasing their chances of evading the predator and enhancing survival. These responses involve a high-energy, anaerobic burst of movement, initiated either from rest or routine swimming (Domenici and Batty, 1994). This burst-swimming behavior can be triggered by the acoustic-lateralis system and the brain stem escape network, including a significant neuron pair called Mauthner cells (M-cells) that process the threat information and rapidly send signals to motor neurons (Eaton et al., 1991;Sillar et al., 2016;Brownstone and Chopek, 2018;Domenici and Hale, 2019). While escape responses can occur in the absence of M-cell firing, the reaction timing and angular speed of the response are significantly slower than if the M cell is stimulated (Hecker et al., 2020). Following activation of this neural network, the body contracts and bends into a C- or S-shape, while typically positioning the head away from the startling stimulus, before rapidly accelerating away from the predator (Yasargil and Diamond, 1968;Sillar et al., 2016). Individuals in social groups must adjust these escape responses to coordinate with their shoal mates, both to limit collisions when moving at high speeds and to maximize predator “confusion” (i.e., the confusion effect, in which predators struggle to focus on a single individual to attack in a rapidly moving and highly coordinate group) (Nadler et al., 2021).

The benefits of sociality to defense become particularly crucial in coral reef environments, which typically have high predation pressure on small-bodied fishes (Hixon and Beets, 1993;Almany and Webster, 2005). Social coral reef fishes develop strategies to avoid predation in these environments, with some species relying on the protection provided by living coral structures and displaying high site fidelity (such as many damselfishes from the Pomacentridae family; Jones et al., 2004). The blue-green Chromis damselfish *Chromis viridis* is one example of a site-attached, live-coral associated fish species, ranging in group sizes composed of just three individuals to larger aggregations comprising hundreds of members (Öhman et al., 1998;Pratchett et al., 2012;Nadler et al., 2014). Several factors contribute to this observed variation in group size. Coral cover and structural complexity are vital factors influencing the distribution and group size of *C. viridis*, with higher coral cover and greater structural complexity offering habitat that can support larger aggregations (Hobbs et al., 2011). Predation risk also plays a role, with higher predation pressure leading to larger group formations as a defensive mechanism (Holbrook et al., 1997). Additionally, competition for resources, such as food and nesting sites, can impact group size, with larger groups forming in areas with abundant resources (Almany, 2004).

In this study, we compared the collective escape behavior and individual escape response among schools consisting of four, eight, and sixteen individuals of the tropical damselfish species *C. viridis*. We hypothesized that schools with fewer individuals would display higher school cohesion and coordination due to a perceived elevated threat level than individuals in larger groups (Killen et al., 2017), resulting in shorter latency times and higher kinematic performance due to greater connectedness between neighbors (Rosenthal et al., 2015) and reduced interactions within the smaller group (Domenici and Batty, 1997). This study seeks to enhance our understanding of the factors influencing escape behavior in animals living in a social context to better understand the fitness consequences for different size groups.

## 2 Materials and Methods

Experimental work was conducted in November to December 2014. The following research was performed with approval from the James Cook University Animal Ethics Committee (approved protocol number A2103), the Great Barrier Reef Marine Park Authority (Permit G13/25909.1), and Queensland Government General Fisheries (Permit 170251).

2.1 Study species, collection, and husbandry

Using monofilament barrier netting, schools of the blue-green Chromis damselfish *C. viridis* (n=336 fish) were captured from reefs in the lagoon adjacent to the Lizard Island Research Station (LIRS) in the northern Great Barrier Reef, Australia (14°40′ 08′′S; 145°27′34′′E) and immediately returned to the flow-through aquarium facilities at LIRS. Schools were maintained in groups of four, eight, and sixteen individuals each (n=36 schools; n=12 per treatment) in 68L tanks (65 cm L × 41 cm W × 40 cm H). Due to the possibility that differences in body size within and among schools could affect performance at both the individual and school level (Morley and Buckel, 2014), body size variation in terms of standard length was minimized both within schools (0.5 cm range from smallest to largest individual in a group) and among schools (mean standard error: 3.32 ± 0.01 cm; range: 2.86-3.70 cm). Fish were fed a body mass specific diet composed of freshly hatched *Artemia* spp. and INVE aquaculture pellets twice daily. To ensure that each school had sufficient time to recover from the stress of collection, all fish were given 7-10 days following collection before experimental testing.

### 2.2 Escape response experimental procedure

Experimental trials were conducted in a laminar flow swim chamber that replicated the natural flow of a coral reef on a calm-weather day (3.2 cm/s; Johansen, 2014). The working section 50 cm L × 40 cm W (filled to 9cm depth), resulting in a total two-dimensional area of 2000 cm^2^, with each fish having 125 cm^2^ (in groups of 16 fish) to 500 cm^2^ (in groups of 4 fish) of space on average to execute behavioral responses. Once each experimental school was placed in the swim chamber, they were acclimated for four hours. Following this acclimation period, the school’s escape response was stimulated using a standardized and reproducible threat protocol, in which a black tapered test tube (2.5 cm diameter × 12 cm length, 37.0 g) was released remotely from 137 cm above the arena using an electromagnet. This stimulus was discharged through a white PVC pipe (to prevent visual detection of the stimulus prior to reaching the water surface) once >50% of the school had gathered in the center of the arena (i.e., more than two body lengths from any arena wall) (Nadler et al., 2021). A piece of fishing line kept this stimulus from striking the experimental arena. Both the tapered shape of the stimulus and the fishing line aided in minimizing any ripples generated when the stimulus first made contact with the water’s surface. Each trial was filmed using a high-speed video camera through a mirror that was angled 45° beneath the transparent swim chamber (240 fps; Casio Exilim HS EX-ZR1000). Between each trial, the swim chamber was drained and refilled with seawater from the LIRS flow-through system.

### 2.3 Behavioral analysis

Video recordings were examined frame by frame using the application Potplayer (v. 1.7.21566) to find crucial points in the individual’s and school’s response to the stimulus. Screenshots of these timepoints were analyzed in ImageJ (v. 1.53n 7). Individual escape performance was evaluated using reaction timing and kinematics, including latency (the interval between the aerial mechanical stimulus first breaking the water’s surface and the fish’s initial head movement), average turning rate (the maximum turning angle, 0, achieved by the fish during stage 1 divided by the time it took to achieve that angle, which serves as a proxy for the fish’s agility through speed of muscle contraction), and distance covered (distance moved in the first 42 ms of the reaction, which is the average time for this species to achieve stages 1 and 2 (Nadler et al., 2021); used as a proxy for swimming speed). Since these traits are influenced by the stimulus distance (the distance between the fish’s center of mass and the stimulus), this trait was also measured and included as a covariate in all analyses (Domenici and Hale, 2019). Non-responders (n=3, 0.8% of all fish included in this study; those fish that did not respond within two seconds of the stimulus) were assigned the maximum measured latency in this study (1003.8 ms), though non-responders were not included in analyses of average turning rate and distance covered. Due to the limits that proximity to wall of the experimental arena can have on kinematic performance (Eaton and Emberley, 1991), the kinematic attributes (average turning rate and distance covered) were only assessed if the fish was >3 cm (i.e., approximately one body length) away from any wall of the experimental arena to ensure that our kinematic data was not impacted by effects imposed by the walls of the arena.

Throughout the response, the school’s cohesion and coordination were measured through nearest neighbor distance and alignment, respectively. Nearest neighbor distance represents the distance between each fish’s center of mass and their most proximal neighbor’s center of mass in the school. Alignment measures the variation in each school’s members orientation with respect to the water’s flow (0°) (Bachelet, 1981). This variation was quantified by using the program Oriana 4 (Kovach, 2011) and determining each school members’ angle then calculating the length of mean circular vector (*r*) of the group, which ranges from 0 (all members are at random angles) to 1 (all angles perfectly aligned). These characteristics were examined at intervals following the stimulus, including 0 ms (representing the school’s cohesiveness and coordination immediately prior to the stimulus), 30 ms (representing the typical time for this species to complete stage 1), and 100 ms (the average time for individuals to complete both stages 1 and 2). Here, time was used as a categorical variable with these time stamps representing different stages of the escape response.

### 2.4 Statistical analyses

R Programming Language (v. 1.3.1093) was used for all statistical analyses (R Development Core Team, 2022). The differences among treatment groups were evaluated using linear mixed-effects models (LMM), using the packages “lme4” (Walker, 2015), “car” (Weisberg, 2019), “MuMIn” (Bartoń, 2023), “emmeans” (Lenth, 2022) and “ggplot2” (Wickham, 2016). Latency, average turning rate, and distance covered were analyzed using LMMs, with group size as a fixed effect, stimulus distance as a covariate, the interaction between group size and stimulus distance, and school identifier as a random effect (so that each individual was nested within their school). School traits (nearest neighbor distance and alignment) were measured with group size, time post-stimulus (0, 30, 100 ms), and their interaction as fixed effects. For the alignment analysis, school identifier was included as a random effect to account for the repeated measures design. Nearest neighbor distance was measured on an individual level, so individual was nested within the school identifier as a random effect. To verify that the assumptions of normality and homogeneity of variance were met for each model, we visually inspected the quantile-quantile and residuals plots and used Shapiro-Wilk and Bartlett tests. To meet these assumptions, all response variables (except average turning rate and alignment) were boxcox transformed using the package “car” (Weisberg, 2019). Best-fit models were identified using the Akaike information criterion (AIC; Hunsicker et al., 2011) model selection. When significant fixed effects were identified, they were further explored using Tukey’s multiple comparison post hoc tests. The R^2^ for all models is also detailed below, including the marginal and conditional R^2^ (R^2^m and R^2^c, respectively), which represent the variance explained by the fixed effects only (R^2^m) and the fixed effects plus the random effects (R^2^c).

## 3 Results

### 3.1 Individual escape performance

For latency, the most complex model was deemed the best-fit through AIC model selection, including group size, stimulus distance, and their interaction as fixed effects. There was no effect of group size on latency (LMM: F_2,35_ = 2.07, p = 0.14, R^2^m = 0.17, R^2^c = 0.30), with all group sizes exhibiting comparable reaction timing. However, there was a non-significant trend in the interaction between group size and stimulus distance (LMM: F_2,307_ = 2.98, p = 0.05; Figure 1), as the relationship between latency and stimulus distance varied with group size. As expected, latency increased (indicating a slower reaction timing) as stimulus distance increased (LMM: F_1,328_ = 52.02, p < 0.0001).

**Figure 1.**
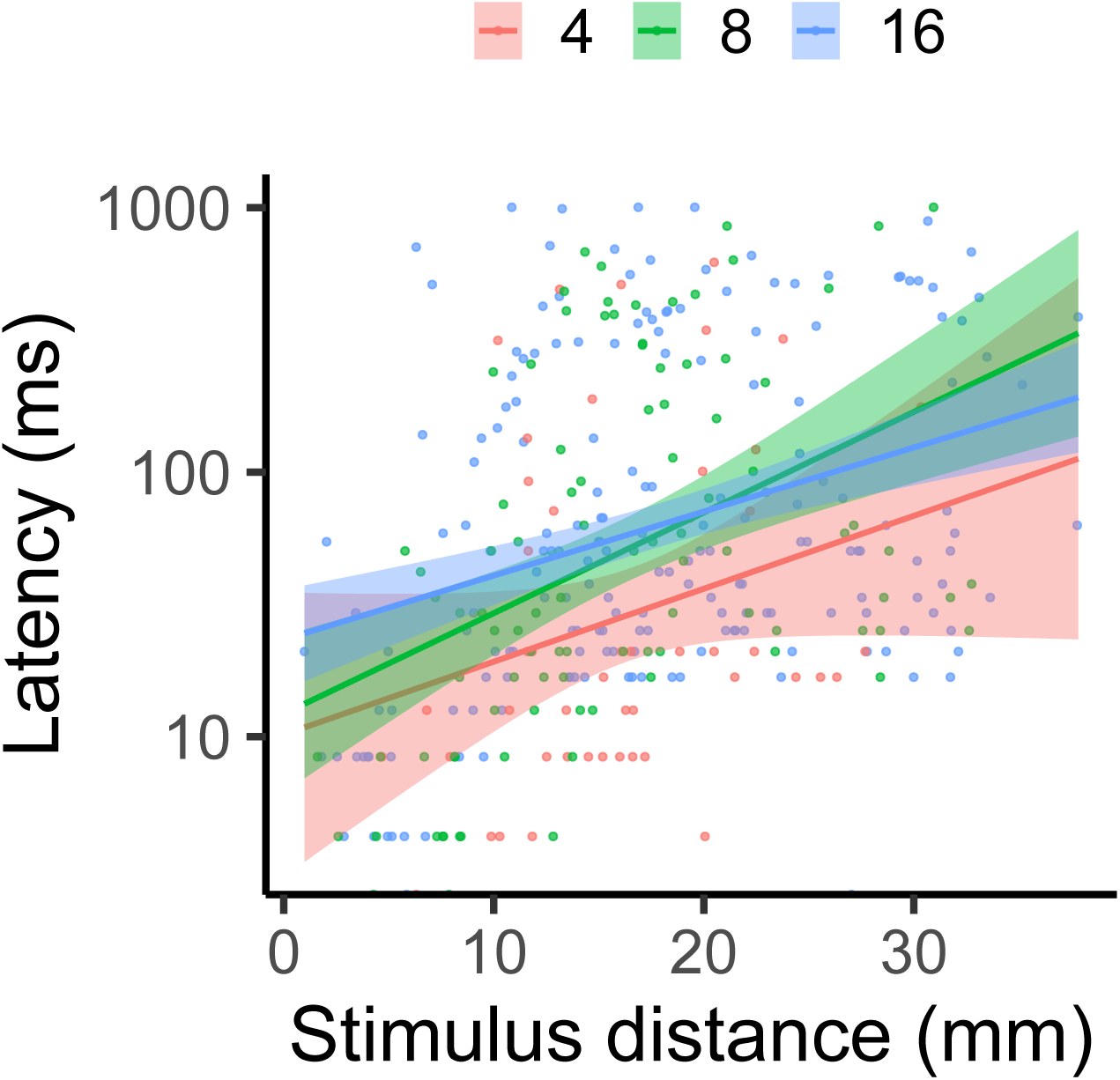
Effect of group size on the latency of escape response in groups composed of four, eight, and sixteen fish, as stimulus distance increases. Each data point represents one individual, with lines indicating a linear regression by treatment and shading representing the 95% CI. The y-axis is logged for visual clarity of trends among treatments.

For average turning rate, the best-fit model was again the most complex model, including group size, stimulus distance, and their interaction. Average turning rate decreased as group size increased (LMM: _F2,34_ = 4.15, p = 0.02, R^2^m= 0.15, R^2^c = 0.33; Figure 2), as individuals from groups of 16 exhibited an approximately 25% lower average turning rate than individuals from groups of four (Tukey’s posthoc test: p_4-16_ = 0.02; all other pairwise comparisons had p > 0.05). Average turning ^rate^ also declined with rising stimulus distance (LMM: F_1,267_ ^= 16.35, p < 0.0001^). There was a significant interaction between group size and stimulus distance (LMM: F_2,267_ ^= 4.33, p = 0.01), due^ to variation in the strength of the relationship between average turning rate and stimulus distance in different group sizes.

**Figure 2.**
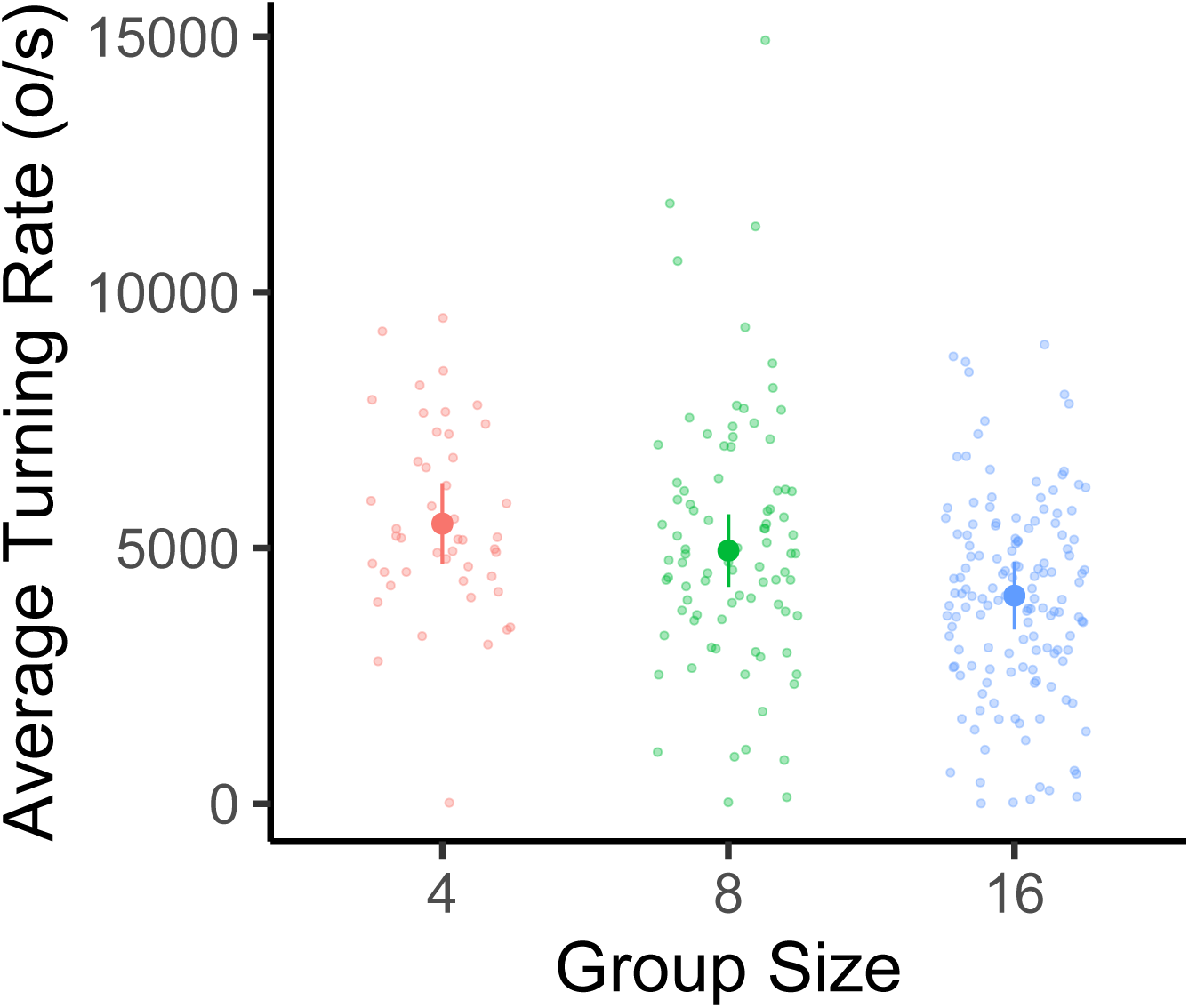
Effect of group size on the average turning rate of the escape response in groups composed of four, eight, and sixteen fish. Large data points represent the estimated marginal mean for each group size (± 95% CI) from linear mixed-effects model analysis (controlling for stimulus distance, with each individual nested within the school with which they were tested). Small data points represent one individual.

The best-fit model for distance covered was the most complex model, including group size, stimulus distance, and their interaction. As with average turning rate, distance covered decreased with group size (LMM: F_2,35_ = 5.51, p = 0.008, R^2^m= 0.18, R^2^c = 0.29; Figure 3), with individuals from groups of sixteen exhibiting a 35% reduction in distance covered when compared to individuals in groups of four (Tukey’s posthoc test: p_4-16_ = 0.005; all other pairwise comparisons had p > 0.05). While distance covered decreased with rising stimulus distance (LMM: F_1,271_ = 36.06, p < 0.0001), the interaction between group size and stimulus distance was not significant (LMM: F_2,263_ = 1.51, p = 0.22).

**Figure 3.**
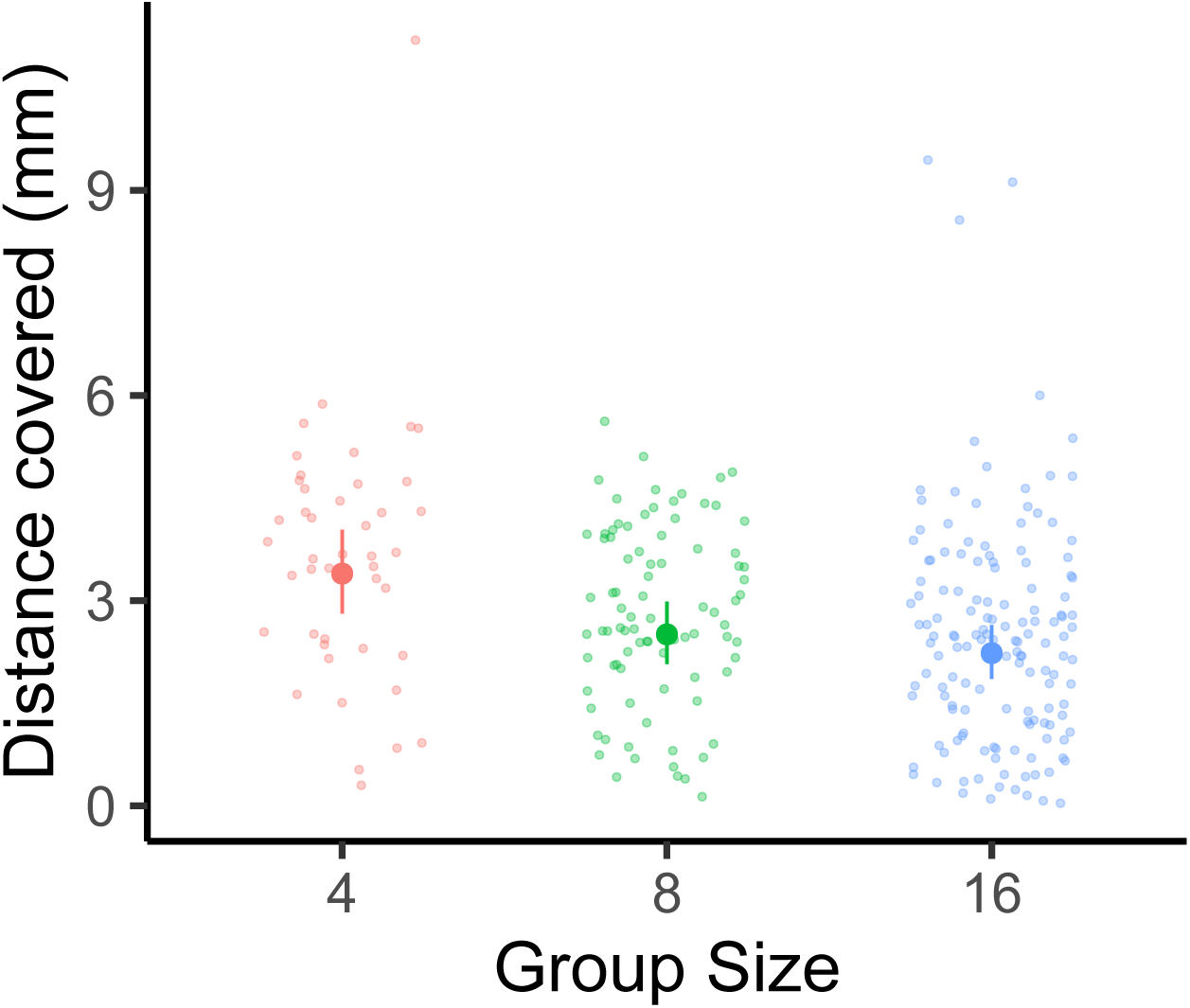
Effect of group size on the distance covered of the escape response in groups composed four, eight, and sixteen fish. Large data points represent the estimated marginal mean for each group size (± 95% CI) from linear mixed-effects model analysis (controlling for stimulus distance and the interaction between stimulus distance and group size, with each individual nested within the school with which they were tested). Small data points represent one individual.

### 3.2 School traits

All school traits were influenced by time after the predator stimulus, while group size impacted how school cohesion (i.e., nearest neighbor distance) changed in the period following the predator stimulus. For nearest neighbor distance, the best-fit model was the most complex model, including group size, time, and their interaction. As group size increased, nearest neighbor distance declined (LMM: F_2,40_ = 14.32, p < 0.0001, R^2^m= 0.09, R^2^c = 0.61; Figure 4), indicating that individuals in larger groups were closer to their nearest neighbor (Tukey’s posthoc test: p_4-8_ = 0.003; p_4-16_ = 0.0003; p_8-16_ > 0.05). Conversely, nearest neighbor distance increased with time post predator stimulus (LMM: F_2,666_ = 10.56, p < 0.0001), as groups decreased cohesion as individuals mounted individual escape responses (Tukey’s posthoc test: p_0-100_ = 0.0004; p_30-100_ = 0.0003). At each time point, individuals in groups of four exhibited an approximately 30-40% higher nearest neighbor distance than groups composed of sixteen fish (Tukey’s posthoc test, 0 ms: p_4-8_ = 0.002; 30 ms: p_4-16_ = 0.0007; ^100 ms: p^_4-16_ < 0.0001). Groups of eight and sixteen were not significantly different at any time point (Tukey’s posthoc test: p > 0.05 for all pairwise comparisons) and the interaction between time and group size was not statistically significant (LMM: F_4,66_ = 1.60, p = 0.17).

**Figure 4.**
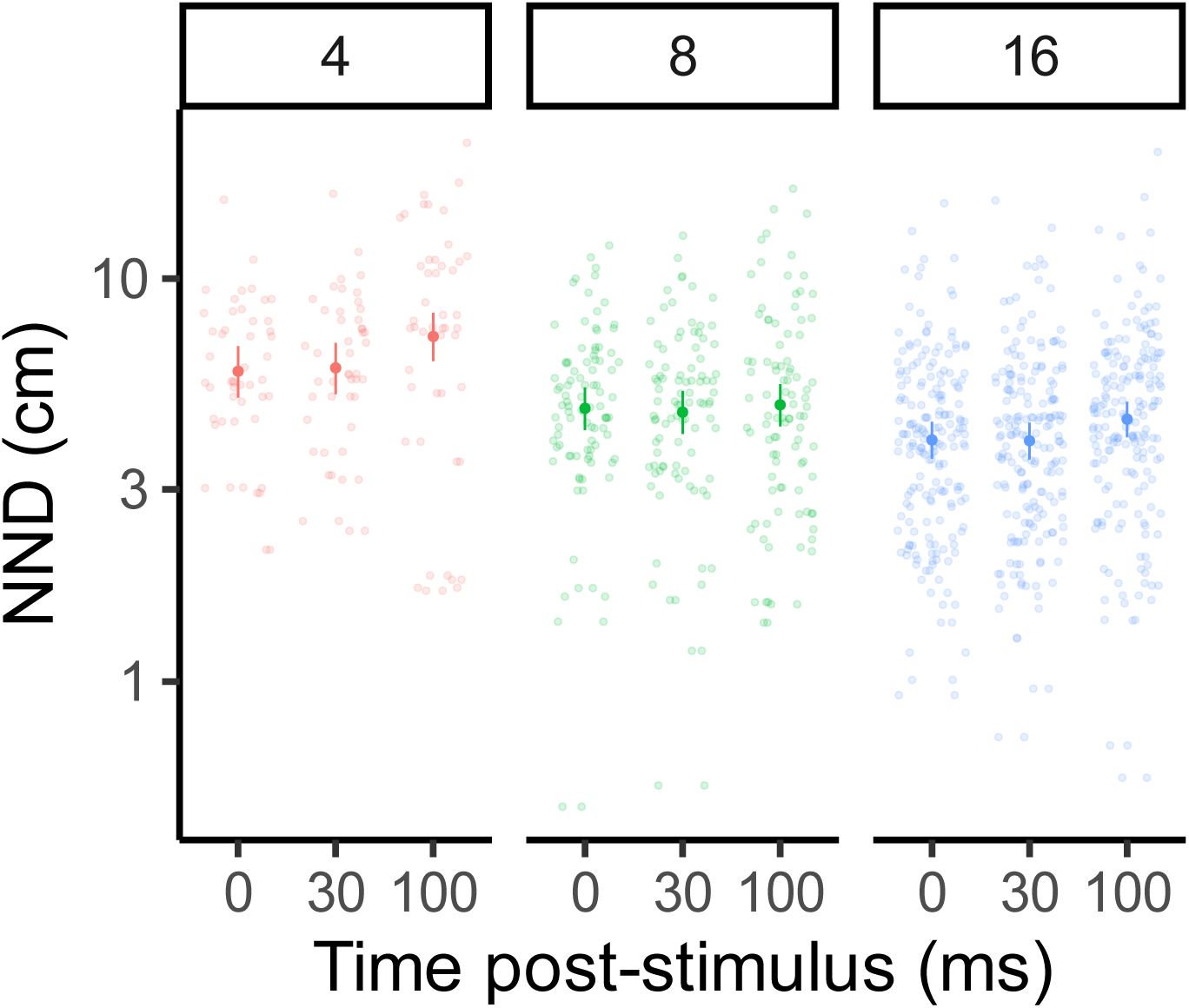
Effect of group size (four, eight, and sixteen fish) on nearest neighbor distance at 0, 30, and 100 ms following a stimulated predator threat. Large data points represent the estimated marginal mean for each group size (± 95% CI) from linear mixed-effects model analysis (controlling for the interaction between stimulus distance and group size, with each individual nested within the school with which they were tested). Small data points represent one individual. Time was included as a categorical fixed effect, as data was only collected at three distinct time points.

For alignment, the best-fit model was the most complex model, including group size, time, and their interaction. While group size did not have a significant effect on alignment (LMM: F_2,33_ = 1.44, p = 0.25, R^2^m= 0.22, R^2^c = 0.64; Figure 5), alignment decreased significantly with time post-simulated predator threat (LMM: F_2,66_ = 21.70, p < 0.0001). Alignment was 17% higher immediately prior to stimulation than it was at 30 and 100 ms after this simulated threat (Tukey’s posthoc test: p_0-100_ < 0.0001; p_0-30_ < 0.0001; p_30-100_ > 0.05). The interaction between group size and time was not significant (LMM: F_4,66_ = 1.98, p = 0.11).

**Figure 5.**
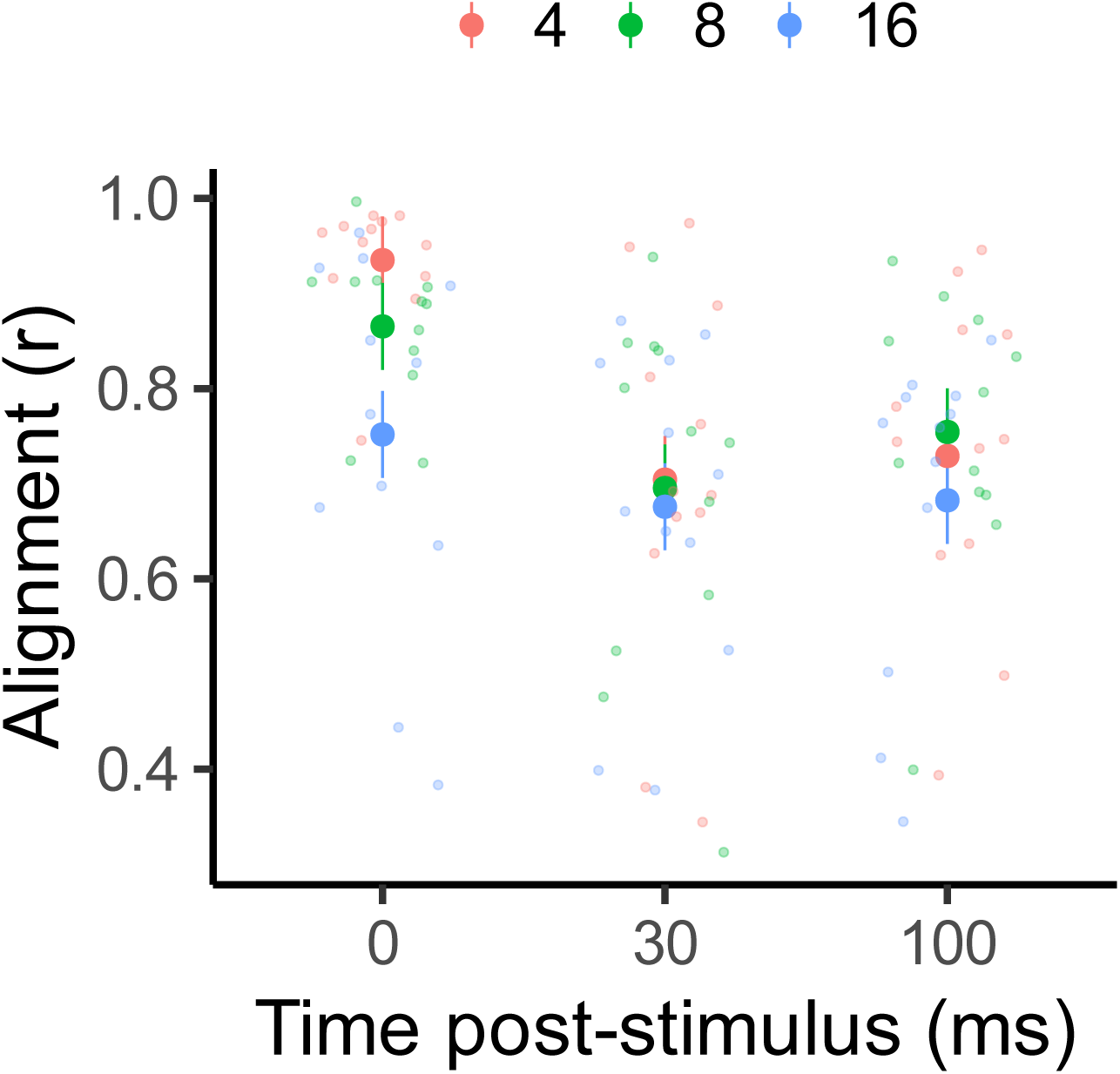
Effect of group size (four, eight, and sixteen fish) on alignment (in terms of length of mean circular vector, *r*) at 0, 30, and 100 ms following a stimulated predator threat. Large data points represent the estimated marginal mean for each group size (± SE) from linear mixed-effects model analysis (controlling for the interaction between stimulus distance and group size, with each individual nested within the school with which they were tested). Small data points represent one individual. Time was included as a categorical fixed effect, as data was only collected at three distinct time points.

## 4 Discussion

The results of this study demonstrate that the escape performance of individual fish is plastic and influenced by social group size. Individuals’ preferred group size can change depending on the context (Magurran et al., 1992;Beauchamp, 2004). For example, group size typically increases in response to a predation cue (i.e., chemical alarm cue) and decreases in response to a food resource (Hoare et al., 2004). While reaction timing (i.e., latency) was consistent across group sizes, kinematic traits, including average turning rate and distance covered, declined as group size increased. The kinematic results confirm those from Domenici & Batty (1997), who measured a higher average turning rate in solitary than schooling fish, similar to the results presented here. While faster performance in solitary fish has been linked to increased escape success (Walker et al., 2005), our results suggest that larger groups exhibit lower performance than small groups. Kinematic performance may have declined with group size due to lower perceived risk of predation due to safety in numbers, higher risk of false alarms, and greater prevalence of collisions with group-mates, which are all more likely to occur in larger schools.

Prey in smaller shoals are more at risk of predation per capita than those in larger shoals (Krause and Godin, 1995), so may need to maintain higher individual kinematic performance to effectively avoid predators. In addition to diluting individual risk of predation, individuals in larger schools benefit from information sharing with group-mates that results in a greater collective intelligence (Papageorgiou and Farine, 2020), including information about predator risk (Magurran and Higham, 1988). As a result, in larger groups, schooling fish may perceive a reduced risk due to safety in numbers through social buffering of stress (Lehtonen and Jaatinen, 2016;Nadler et al., 2016;Culbert et al., 2019;Yusishen et al., 2020), and hence reduce investment in kinematic performance. For example, in the gregarious cichlid *Neolamprologus pulcher*, individuals mount a reduced behavioral (activity) and physiological (cortisol) stress response to a predation-like stressor (i.e., air exposure) when they recovered with group-mates than alone (Culbert et al., 2019). Conversely, larger groups may experience a higher frequency of false alarms, where individuals mistakenly initiate rapid defensive maneuvers, which is likely more prevalent in larger group sizes (Gray and Webster, 2023). Similarly, due to smaller nearest neighbor distances, fish in larger schools may be more at risk of collisions with group-mates, so fish may reduce their escape speed to limit risk of colliding other rapidly moving fish (Herbert-Read, 2016). The M-cell response is highly plastic, with evidence suggesting that responses in the absence of M-cell firing result in reduced kinematic performance (Hecker et al., 2020), as seen here in fish from larger group sizes. Any factor that increases the threshold of threat necessary to initiate a rapid, M-cell driven escape response, such as perceived safety, frequency of false alarms, or collision risk, could act to reduce each individual’s kinematic performance.

In contrast to past studies that compared solitary and schooling fishes (Webb, 1980;Domenici and Batty, 1997), we found no change in reaction timing (i.e., latency) as group size increased. Social context could modulate latency for a range of reasons. A faster latency could be expected as predator detection increases with larger group sizes or responsiveness increases with greater socially transmitted information in a larger group (Treherne and Foster, 1980;Godin and Abrahams, 1988). Conversely, a slower latency may occur if schooling fish are distracted by unfamiliar group mates (Nadler et al., 2021) or if fish are attempting to more accurately determine the most effective escape trajectory based on socially-transmitted information (Domenici and Batty, 1994). While Domenici and Batty (1997) measured a slower latency in schooling versus solitary fish, Webb (1980) found that several schooling fishes either reacted more quickly or the same as solitary counterparts. Here, we found no change in latency with group size, which could stem from several sources. Unlike Domenici and Batty (1997) and Webb (1980), we compared latency across a group-size gradient, rather than a solitary versus social condition. Further, we used different species and predator stimuli (mechanical, versus sound as in Domenici and Batty 1997 or electric shock as in Webb 1980), which may affect the initiation of the escape response.

Group cohesion increased with group size, suggesting that school performance may have increased despite reduced individual performance. Partridge (1980) found a similar effect in schooling minnows (*Phoxinus phoxinus*), in which nearest neighbor distance decreased by more than half when the group size increase from two to four individuals. In the study presented here, as the size of the arena was consistent across treatments, the density in the experimental arena would also have increased with group size. In schooling herring (*Clupea harengus*), Rieucau et al. (2014b) found that higher density schools execute more cohesive collective escape responses than lower density schools, potentially due to faster propagation of social information between neighboring fish than under lower density conditions. Further, in larger groups, individuals may join smaller sub-groupings to promote cohesion and swimming performance among its members, with the larger group acting according to the consensus of its sub-groups (Hemelrijk and Hildenbrandt, 2012;Kao and Couzin, 2019). While density of group-mates was inevitably higher in the larger groups, all fish had at least 125 cm^2^ on average to execute their responses (see details in Materials and Methods section). Thus, space limitation was likely not the driving factor altering kinematic performance. However, research into the space requirements necessary for ecologically relevant behavioral measurements in social fishes would be useful to confirm space limitation did not influence these findings.

To successfully survive a predator’s attack, prey must effectively initiate rapid defensive behavior (Walker et al., 2005;McCormick and Allan, 2017). Synchronization was traditionally thought to be an essential component of a successful collective antipredator response to minimize the oddity effect (Landeau and Terborgh, 1986;Conradt and Roper, 2000) and maximize predator confusion caused by the coordinated movements of its group-living prey (Marras et al., 2012). In some contexts, confusion may be generated more effectively when the prey’s response is random than if the group remains highly coordinated (Ruxton et al., 2007). A study on the same species, *C. viridis*, observed a similar effect, in which the group’s cohesion and coordination declined in the 30-100 ms following the predator’s attack (Nadler et al., 2021). Similar reductions in group cohesion and coordination immediately following the predator attack have been seen in other fishes as well. Romenskyy et al. (2020) found that schools of the Pacific blue-eye (*Pseudomugil signifier*) responded to an attack by the predatory flat-headed gudgeon (*Philypnodon grandiceps*) through a so-called “flash expansion”, a rapid increase in the school’s volume in three dimensions but recovered quickly to baseline levels.

While laboratory studies are useful for controlling variation among treatments, natural conditions are rarely fully replicated in a laboratory setting (Campbell et al., 2009). Therefore, field studies are crucial to validate the functional consequences of these behavioral changes to prey survival of predator attacks (McCormick et al., 2018). Freely formed group sizes are highly context dependent, with fish establishing schools of varying sizes in response to food or alarm cues (Hoare et al., 2004). However, while there is risk that cohesion of larger groups was constrained by the size of the experimental arena, *C. viridis* are typically found in large aggregations on coral reefs (Hobbs et al., 2011), so a larger group size will often result in the higher density conditions in this species . In follow-up studies, allowing schooling fishes to freely form various school sizes may further validate the results found in the present study.

In conclusion, we found that kinematic performance declines in fish in larger groups, though larger group sizes maintain stronger group cohesion than smaller group sizes before, during, and after a predator attack. Though we typically associate a faster escape performance with a greater chance of survival of predator attacks (Walker et al., 2005), these findings suggest that fish in larger groups may be able to save energy on mounting energetically-costly behavioral responses while evading predator attacks due to a greater perceived safety in numbers (Lehtonen and Jaatinen, 2016). The greater capacity for predator vigilance in larger group sizes comes with the tradeoff of greater competition for limited resources, such as food and habitat (Gil et al., 2017), which theoretically results in an optimal group size that balances these costs against potential benefits (Brown, 1982).

## Supporting information

Bacchus et al._R Markdown

Alignment data

Fast start data (no na's)

Fast start data (all)

Nearest neighbor distance data

## Author Contributions

L.E.N., P.D., S.S.K. and M.I.M. designed the concepts and methodology for the study. L.E.N. and S.S.K. conducted the experiment. M.D.B. and L.E.N. conducted the video and statistical analyses. M.D.B., P.D. and L.E.N interpreted the data. M.D.B. and L.E.N. drafted the manuscript. All authors edited the manuscript and gave approval for publication.

## Acknowledgments

We thank Eva McClure, the staff at the Lizard Island Research Station (a facility of the Australian Museum), Ross Barrett and Stephen Brown for logistical support. In addition, thank you to Professors David Kerstetter and Omar Eldakar for serving on the committee for M.D.B.’s Master thesis, which included this study. Lastly, many thanks to the undergraduate students that assisted with video analysis at various stages, including Robby Spekis, Alianna Jones, Delaney Farrell, James Puentes, and Raghavi Vuppala.

