## Supplementary material for "Kinematic performance declines as group size increases during escape responses in a schooling coral reef fish": Bacchus et al._R Markdown

2023-09-12

Escaping predation is essential for species survival, but prey must effectively match their response to the perceived threat imposed by a predator. For social animals, one mechanism to reduce risk of predation is living in larger group sizes, which dilutes each individual's risk of capture. When a predator attacks, individuals from a range of taxa (e.g., fishes, sharks, amphibians) use the rapid, anaerobically-fueled burst swimming behavior, known as the fast-start response, to evade the attack. Here, using the schooling coral reef damselfish Chromis viridis, we assess if there is an optimal group size that maximizes both individual escape response as well as group cohesion and coordination following a simulated predator attack, comparing schools composed of four, eight, and sixteen fish. We found that fish in various group sizes exhibited no difference in their reaction timing to a simulated predator attack (i.e., latency), but larger groups exhibited a slower kinematic response (i.e., lower average turning rate and shorter distance covered during the escape response), potentially because larger groups perceived the predator attack as less risky due to safety in numbers. Both school cohesion and coordination (as measured through alignment and nearest neighbor distance, respectively) declined in the 100ms after the predator's attack. While there was no impact of group size on alignment, larger group sizes exhibited closer nearest neighbor distance at all time stamps. This study highlights that larger group sizes may allow individuals to save energy on costly behavioral responses to avoid predators, potentially through a greater threshold of threat necessary to trigger a rapid escape response.

```
#LIBRARIES
library(lme4)
library(MuMIn)
library(emmeans)
library(car)
library(ggplot2)
```

### LATENCY

Latency, or reaction timing, was measured as the time interval (in milliseconds) between the aerial mechanical stimulus first breaking the water's surface and the fish's initial head movement. Non-responders (n=3, 0.8% of all fish included in this study; those fish that did not respond within two seconds of the stimulus) were assigned the maximum measured latency in this study (1003.8 ms).

```
fs_all <- read.csv("/Users/laurennadler/Desktop/MyRData/Monica_groupsize_faststart.csv")
fs_all$Group.Size = as.factor(fs_all$Group.Size)
lat.hist <- ggplot(fs_all, aes(x=Latency, color = Group.Size, fill = Group.Size)) +
  geom_histogram(position = "dodge", alpha = 0.5, bins = 6) +
  theme_classic(base_family='Arial', base_size = 28) +
  theme(legend.title=element_blank()) +
  theme(legend.position="top")
lat.hist
```

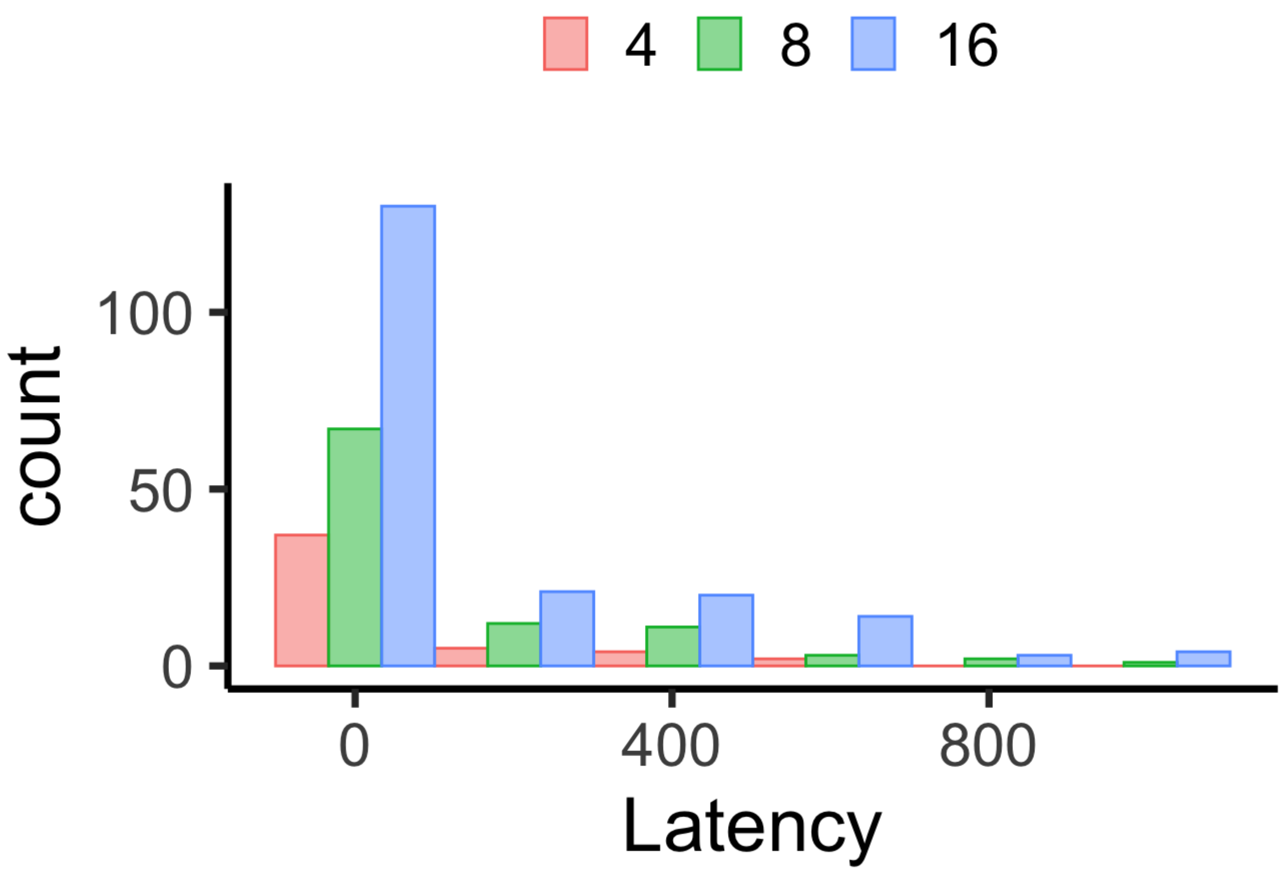


```
#latency model
allLat.lmer = lmer((Latency + 0.001)~ Group.Size*Stimulus.distance + (1|Video), data=fs_
all, na.action = "na.omit")
plot(allLat.lmer) #highly heteroscedastic
```

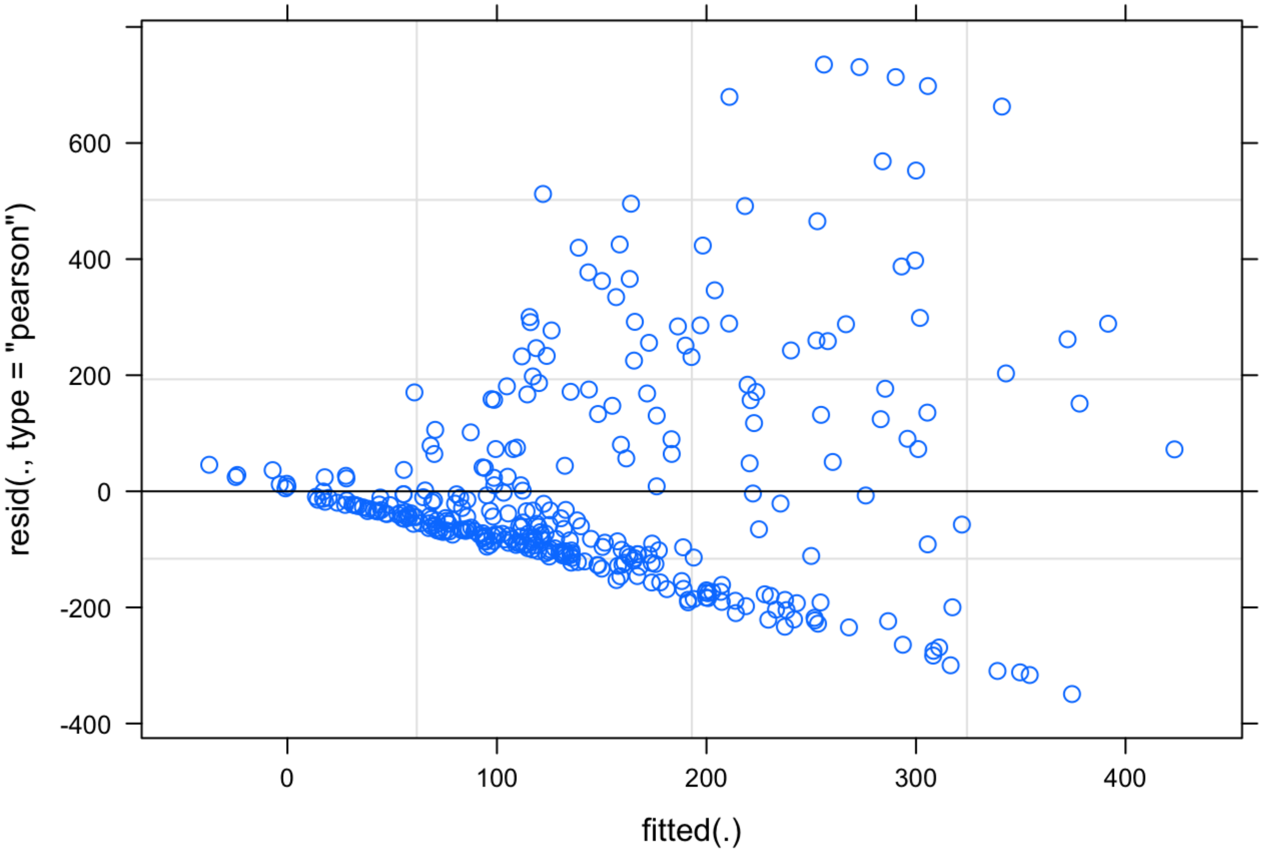


```
qqnorm(resid(allLat.lmer))
qqline(resid(allLat.lmer))
```

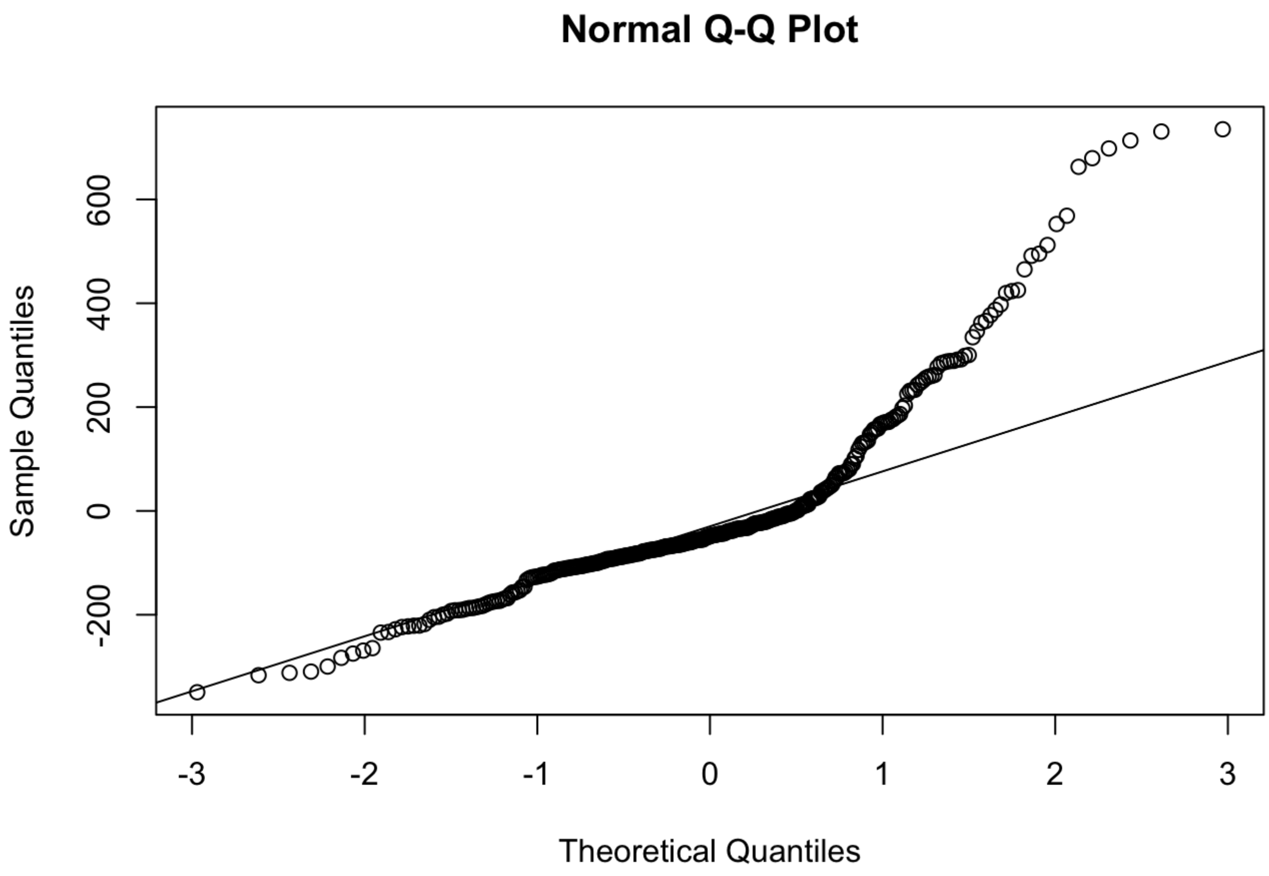


```
shapiro.test(resid(allLat.lmer)) #p=9.752e-09 so non normal
```

```
##
##  Shapiro-Wilk normality test
##
## data:  resid(allLat.lmer)
## W = 0.85535, p-value < 2.2e-16
```

```
bartlett.test(resid(allLat.lmer), fs_all$Group.Size) #p=0.03 so heteroscedastic
```

```
##
##  Bartlett test of homogeneity of variances
##
## data:  resid(allLat.lmer) and fs_all$Group.Size
## Bartlett's K-squared = 16.736, df = 2, p-value = 0.0002322
```

```
allLat.lmer.bc <- powerTransform(allLat.lmer, family="bcPower") #used boxcox transformat
ion due to violation of assumptions of normality and homoscedasticity
summary(allLat.lmer.bc) #both significant
```

```
## bcPower Transformation to Normality
##      Est Power Rounded Pwr Wald Lwr Bnd Wald Upr Bnd
## [1,]    0.1477        0.15       0.1116       0.1837
##
## Likelihood ratio test that transformation parameter is equal to 0
##  (log transformation)
##                              LRT df       pval
## LR test, lambda = (0) 96.44479  1 < 2.22e-16
##
## Likelihood ratio test that no transformation is needed
##                             LRT df       pval
## LR test, lambda = (1) 702.557  1 < 2.22e-16
```

```
allLat.lmer.bc$roundlam
```

```
## [1] 0.1476753
```

```
allLat.bc.lmer = lmer(bcPower((Latency+0.001), allLat.lmer.bc$roundlam) ~  Group.Size*St
imulus.distance + (1|Video), data=fs_all, na.action = "na.omit")
plot(allLat.bc.lmer)
```

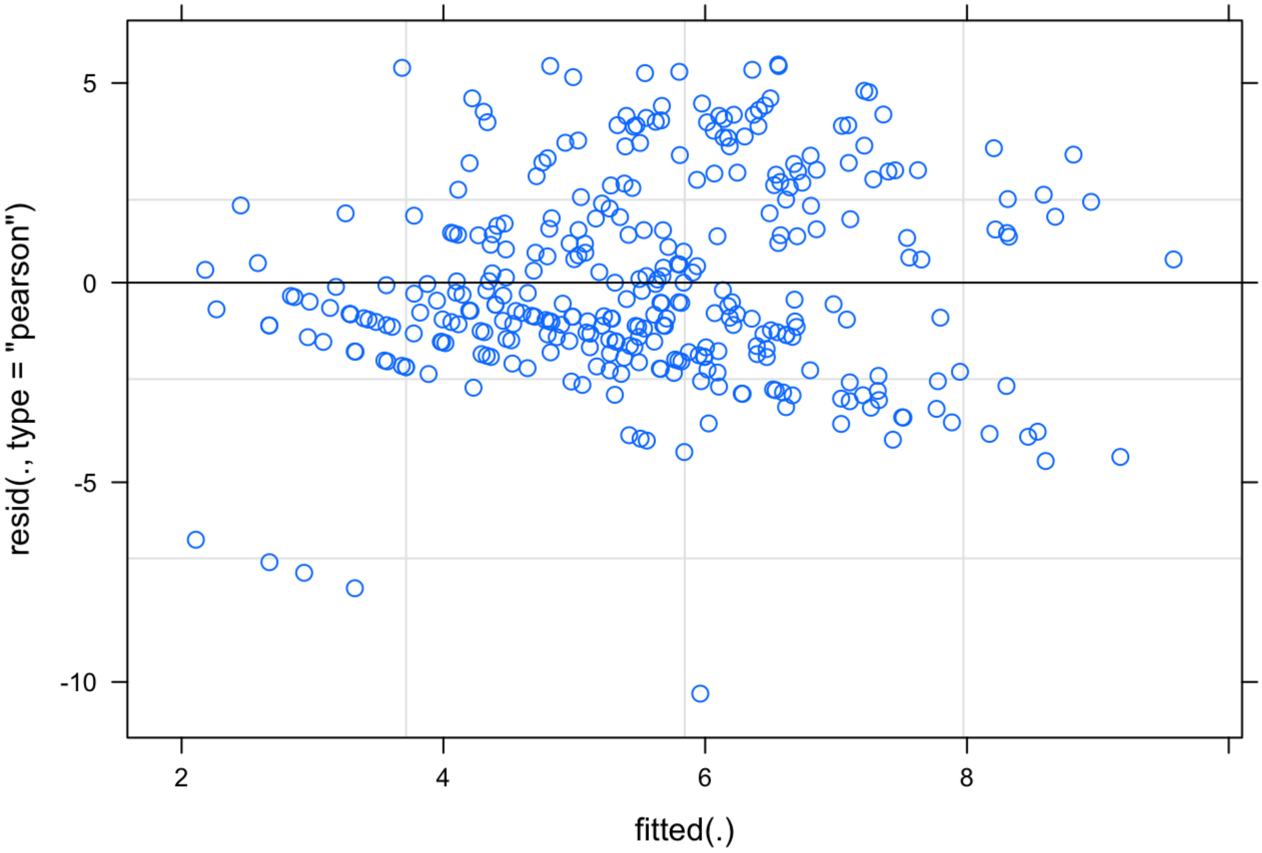

```
qqnorm(resid(allLat.bc.lmer))
qqline(resid(allLat.bc.lmer))
```

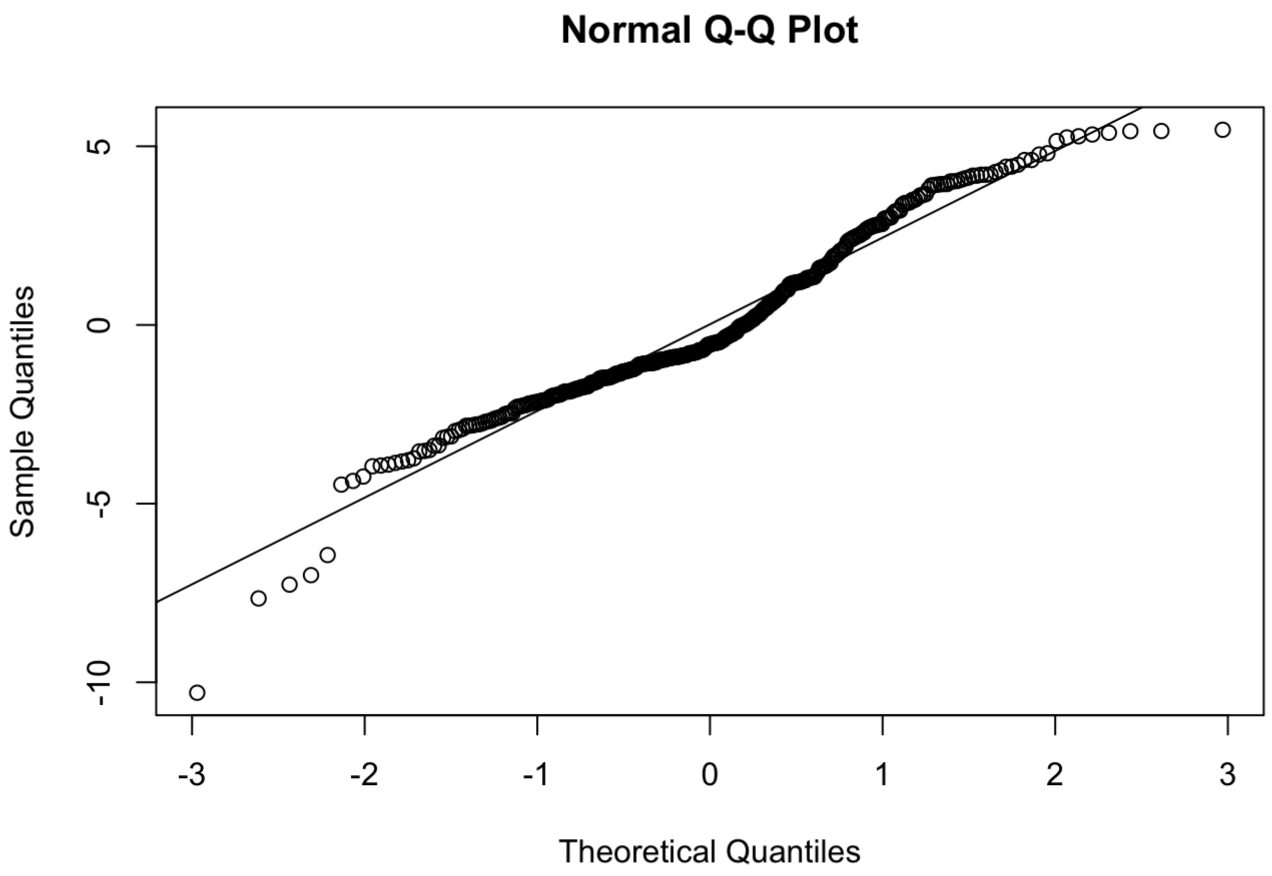


```
shapiro.test(resid(allLat.bc.lmer)) #still p < 0.05 but visual inspection of residuals p
lot dramatically improved by boxcox transformation so continuing with lmer with boxcox t
ransformation
```

```
##
##  Shapiro-Wilk normality test
##
## data:  resid(allLat.bc.lmer)
## W = 0.9668, p-value = 6.035e-07
```

```
bartlett.test(resid(allLat.bc.lmer), fs_all$Group.Size)
```

```
##
##  Bartlett test of homogeneity of variances
##
## data:  resid(allLat.bc.lmer) and fs_all$Group.Size
## Bartlett's K-squared = 0.55796, df = 2, p-value = 0.7566
```

```
allLat.bc.lmer2 = lmer(bcPower((Latency+0.001), allLat.lmer.bc$roundlam) ~  Group.Size+S
timulus.distance + (1|Video), data=fs_all, na.action = "na.omit")
anova(allLat.bc.lmer,allLat.bc.lmer2) #more complex model has a lower AIC so maintaining
allLat.bc.lmer
```

```
## Data: fs_all
## Models:
## allLat.bc.lmer2: bcPower((Latency + 0.001), allLat.lmer.bc$roundlam) ~ Group.Size + S
timulus.distance + (1 | Video)
## allLat.bc.lmer: bcPower((Latency + 0.001), allLat.lmer.bc$roundlam) ~ Group.Size * St
imulus.distance + (1 | Video)
##                 npar    AIC    BIC  logLik deviance  Chisq Df Pr(>Chisq)
## allLat.bc.lmer2    6 1641.8 1664.7 -814.89   1629.8
## allLat.bc.lmer     8 1640.1 1670.6 -812.05   1624.1 5.6848  2    0.05829 .
## ---
## Signif. codes:  0 '***' 0.001 '**' 0.01 '*' 0.05 '.' 0.1 ' ' 1
```

```
summary(allLat.bc.lmer)
```

```
## Linear mixed model fit by REML ['lmerMod']
## Formula: bcPower((Latency + 0.001), allLat.lmer.bc$roundlam) ~ Group.Size *
##     Stimulus.distance + (1 | Video)
##    Data: fs_all
##
## REML criterion at convergence: 1637.6
##
## Scaled residuals:
##     Min      1Q  Median      3Q     Max
## -3.9531 -0.6206 -0.2099  0.6365  2.0979
##
## Random effects:
##  Groups   Name        Variance Std.Dev.
##  Video    (Intercept) 1.233    1.110
##  Residual             6.779    2.604
## Number of obs: 336, groups:  Video, 36
##
## Fixed effects:
##                                Estimate Std. Error t value
## (Intercept)                     1.29852    1.27706   1.017
## Group.Size8                     0.63556    1.48855   0.427
## Group.Size16                    2.63318    1.39083   1.893
## Stimulus.distance               0.19288    0.07251   2.660
## Group.Size8:Stimulus.distance   0.02542    0.08317   0.306
## Group.Size16:Stimulus.distance -0.08360    0.07602  -1.100
##
## Correlation of Fixed Effects:
##             (Intr) Grp.S8 Gr.S16 Stmls. G.S8:S
## Group.Size8 -0.858
## Group.Siz16 -0.918  0.788
## Stmls.dstnc -0.922  0.791  0.847
## Grp.Sz8:St.  0.804 -0.901 -0.738 -0.872
## Grp.Sz16:S.  0.880 -0.755 -0.895 -0.954  0.832
```

```
Anova(allLat.bc.lmer, test="F")
```

```
## Analysis of Deviance Table (Type II Wald F tests with Kenward-Roger df)
##
## Response: bcPower((Latency + 0.001), allLat.lmer.bc$roundlam)
##                                   F Df Df.res    Pr(>F)
## Group.Size                   2.0670  2  35.42   0.14156
## Stimulus.distance           52.0240  1 328.29 3.812e-12 ***
## Group.Size:Stimulus.distance  2.9768  2 306.85   0.05243 .
## ---
## Signif. codes:  0 '***' 0.001 '**' 0.01 '*' 0.05 '.' 0.1 ' ' 1
```

```
r.squaredGLMM(allLat.bc.lmer)
```

```
##              R2m       R2c
## [1,] 0.1701426 0.2978518
```

```
latency.plot <- ggplot(fs_all, aes(x=Stimulus.distance, y=Latency, colour=Group.Size)) +
  geom_point(aes(color = Group.Size), alpha = 0.7, size=1) +
  geom_smooth(method="glm", fullrange=TRUE, aes(fill = Group.Size)) +
  xlab(bquote('Stimulus distance (mm)')) + ylab(bquote('Latency (ms)')) +
  theme_classic(base_family='Arial', base_size = 28) +
  theme(legend.title=element_blank()) +
  scale_y_log10() +
  theme(legend.position = "top")
latency.plot
```

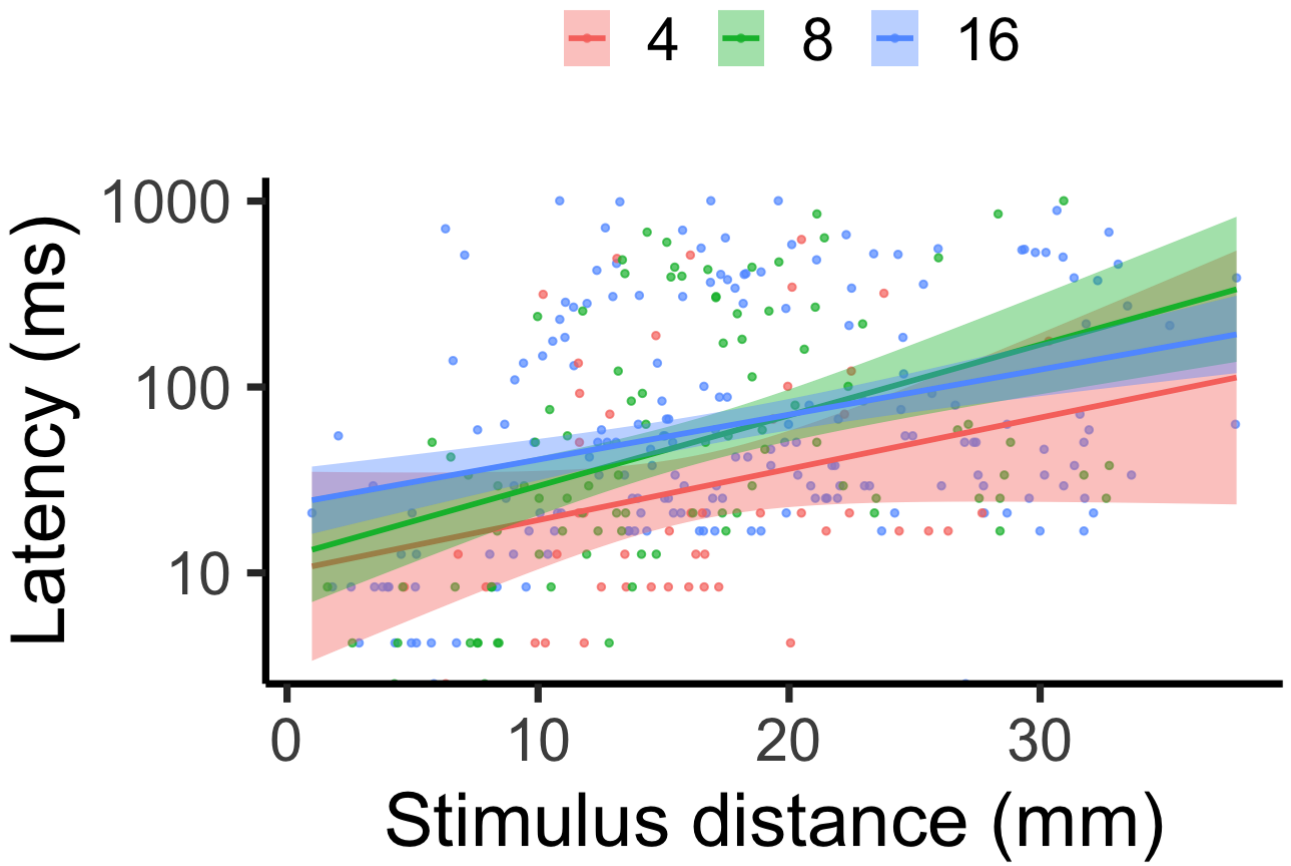


### AVERAGE TURNING RATE & DISTANCE COVERED

We quantified fast-start characteristics associated with the initial unilateral body bend following stimulation (i.e., stage 1 of the fast-start) and the subsequent contralateral body bend (i.e., stage 2 of the fast-start). Individual fast-start kinematic performance was characterized through average turning rate (AVT, the maximum turning angle achieved by the fish during stage 1 divided by the time it took to achieve that angle, which serves as a proxy for the response's agility through speed of muscle contraction), and distance covered (DC; distance moved in the first 42 ms of the reaction, which is the average time for this species to achieve stages 1 and 2; used as a proxy for swimming speed and acceleration). Since these traits are influenced by the stimulus distance (the distance between the fish's center of mass and the stimulus), this trait was also measured and included as a covariate in all analyses (including latency above).

```
#avt - NAs removed (ie - fish too close to wall and non-responders)
fs_noNA <- read.csv("/Users/laurennadler/Desktop/MyRData/Monica_groupsize_faststart no n
as.csv")
fs_noNA$Group.Size = as.factor(fs_noNA$Group.Size)
avt.lmer = lmer(Average.Turning.Rate.per.s ~ Group.Size*Stimulus.distance + (1|Video), d
ata=fs_noNA, na.action = "na.omit")
plot(avt.lmer)
```

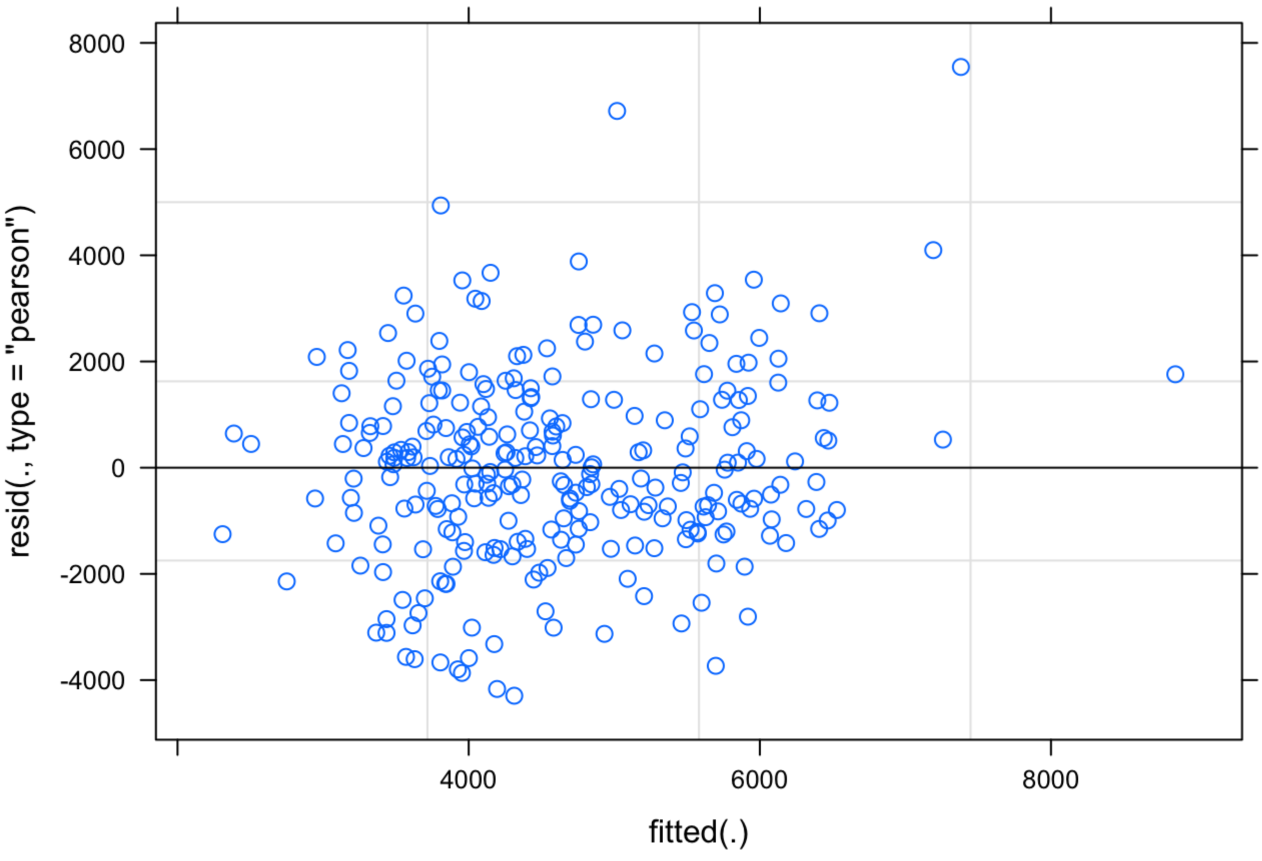


```
qqnorm(resid(avt.lmer))
qqline(resid(avt.lmer))
```

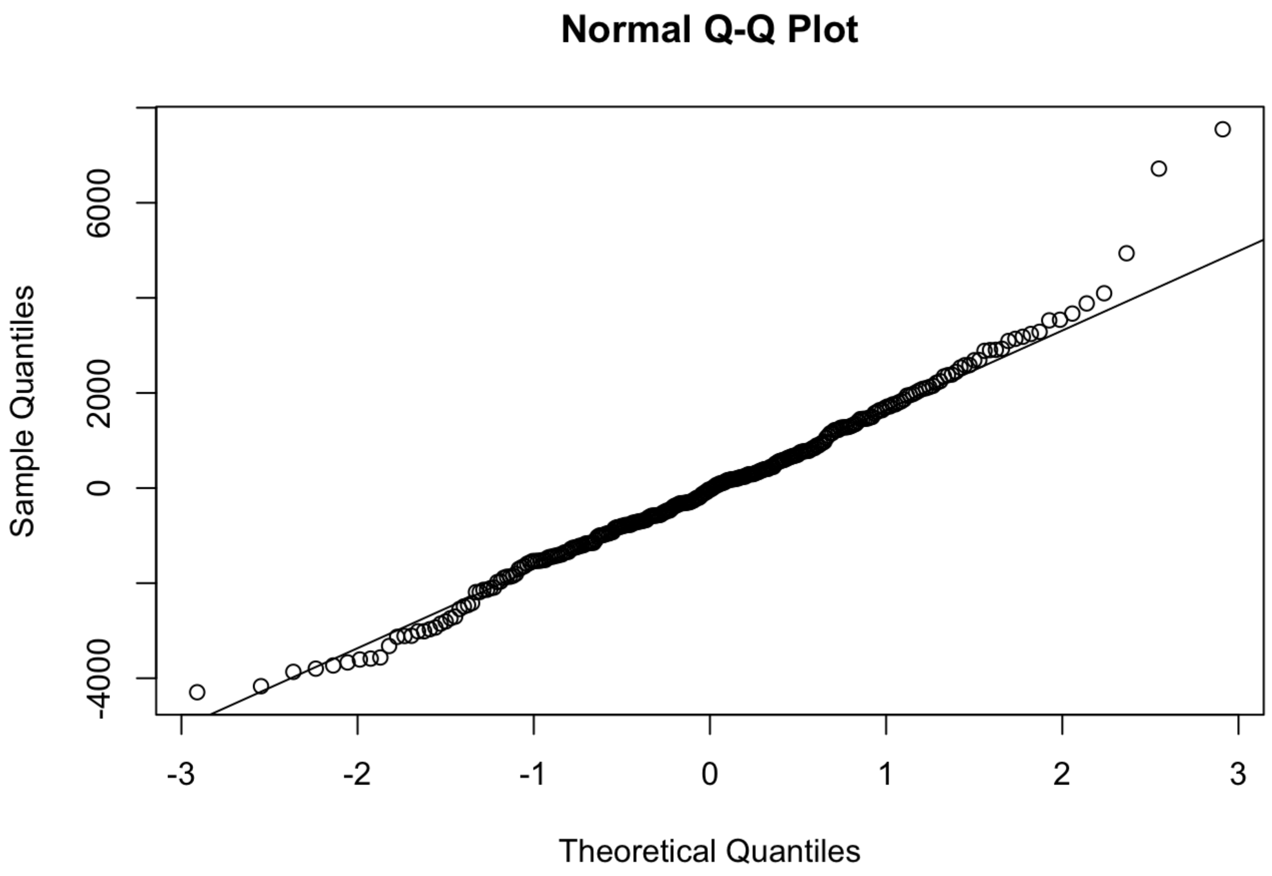


```
shapiro.test(resid(avt.lmer)) #p = 0.003 suggesting non normal but disregarding because
the qqplot looks fine
```

```
##
##  Shapiro-Wilk normality test
##
## data:  resid(avt.lmer)
## W = 0.98343, p-value = 0.002679
```

```
bartlett.test(resid(avt.lmer), fs_noNA$Group.Size)
```

```
##
##  Bartlett test of homogeneity of variances
##
## data:  resid(avt.lmer) and fs_noNA$Group.Size
## Bartlett's K-squared = 2.8727, df = 2, p-value = 0.2378
```

```
avt.lmer2 = lmer(Average.Turning.Rate.per.s ~ Group.Size+Stimulus.distance + (1|Video),
data=fs_noNA, na.action = "na.omit")
anova(avt.lmer,avt.lmer2) #suggests that the more complex lmer better
```

```
## Data: fs_noNA
## Models:
## avt.lmer2: Average.Turning.Rate.per.s ~ Group.Size + Stimulus.distance + (1 | Video)
## avt.lmer: Average.Turning.Rate.per.s ~ Group.Size * Stimulus.distance + (1 | Video)
##            npar    AIC    BIC  logLik deviance Chisq Df Pr(>Chisq)
## avt.lmer2     6 5014.5 5036.2 -2501.2   5002.5
## avt.lmer      8 5010.5 5039.5 -2497.3   4994.5 7.961  2    0.01868 *
## ---
## Signif. codes:  0 '***' 0.001 '**' 0.01 '*' 0.05 '.' 0.1 ' ' 1
```

```
summary(avt.lmer)
```

```
## Linear mixed model fit by REML ['lmerMod']
## Formula: Average.Turning.Rate.per.s ~ Group.Size * Stimulus.distance +
##     (1 | Video)
##    Data: fs_noNA
##
## REML criterion at convergence: 4926.9
##
## Scaled residuals:
##     Min      1Q  Median      3Q     Max
## -2.2855 -0.6135 -0.0182  0.5859  4.0162
##
## Random effects:
##  Groups   Name        Variance Std.Dev.
##  Video    (Intercept)  933892   966.4
##  Residual             3531008  1879.1
## Number of obs: 277, groups:  Video, 36
##
## Fixed effects:
##                                 Estimate Std. Error t value
## (Intercept)                     6183.148   1037.116   5.962
## Group.Size8                     1449.918   1218.881   1.190
## Group.Size16                   -1442.735   1156.860  -1.247
## Stimulus.distance                -43.418     59.125  -0.734
## Group.Size8:Stimulus.distance   -121.395     68.846  -1.763
## Group.Size16:Stimulus.distance     2.114     63.699   0.033
##
## Correlation of Fixed Effects:
##             (Intr) Grp.S8 Gr.S16 Stmls. G.S8:S
## Group.Size8 -0.851
## Group.Siz16 -0.896  0.763
## Stmls.dstnc -0.925  0.787  0.829
## Grp.Sz8:St.  0.795 -0.903 -0.712 -0.859
## Grp.Sz16:S.  0.859 -0.731 -0.899 -0.928  0.797
```

```
Anova(avt.lmer, test="F")
```

```
## Analysis of Deviance Table (Type II Wald F tests with Kenward-Roger df)
##
## Response: Average.Turning.Rate.per.s
##                                    F Df  Df.res   Pr(>F)
## Group.Size                    4.1548  2  33.905  0.02433 *
## Stimulus.distance            16.3507  1 267.466 6.89e-05 ***
## Group.Size:Stimulus.distance  4.3305  2 267.378  0.01410 *
## ---
## Signif. codes:  0 '***' 0.001 '**' 0.01 '*' 0.05 '.' 0.1 ' ' 1
```

```
r.squaredGLMM(avt.lmer)
```

```
##            R2m       R2c
## [1,] 0.1484504 0.3265631
```

```
emmeans(avt.lmer, pairwise~Group.Size, adjust = c("Tukey"))
```

```
## $emmeans
##  Group.Size emmean  SE   df lower.CL upper.CL
##  4            5476 394 52.9     4686     6266
##  8            4949 347 32.1     4242     5656
##  16           4068 321 23.6     3406     4730
##
## Degrees-of-freedom method: kenward-roger
## Confidence level used: 0.95
##
## $contrasts
##  contrast                   estimate  SE   df t.ratio p.value
##  Group.Size4 - Group.Size8       527 525 42.0   1.003  0.5788
##  Group.Size4 - Group.Size16     1408 508 37.0   2.774  0.0229
##  Group.Size8 - Group.Size16      882 473 27.7   1.866  0.1677
##
## Degrees-of-freedom method: kenward-roger
## P value adjustment: tukey method for comparing a family of 3 estimates
```

```
avt.plot <- ggplot(fs_noNA, aes(x=Stimulus.distance, y=Average.Turning.Rate.per.s, colou
r=Group.Size)) +
  geom_point(aes(color = Group.Size), alpha = 0.7, size=1) +
  geom_smooth(method="glm", fullrange=TRUE, aes(fill = Group.Size)) +
  xlab(bquote('Stimulus distance (cm)')) + ylab(bquote('ATR (o/s)')) +
  theme_classic(base_family='Arial', base_size = 20) +
  theme(legend.title=element_blank()) +
  theme(legend.position = "top")
avt.plot
```

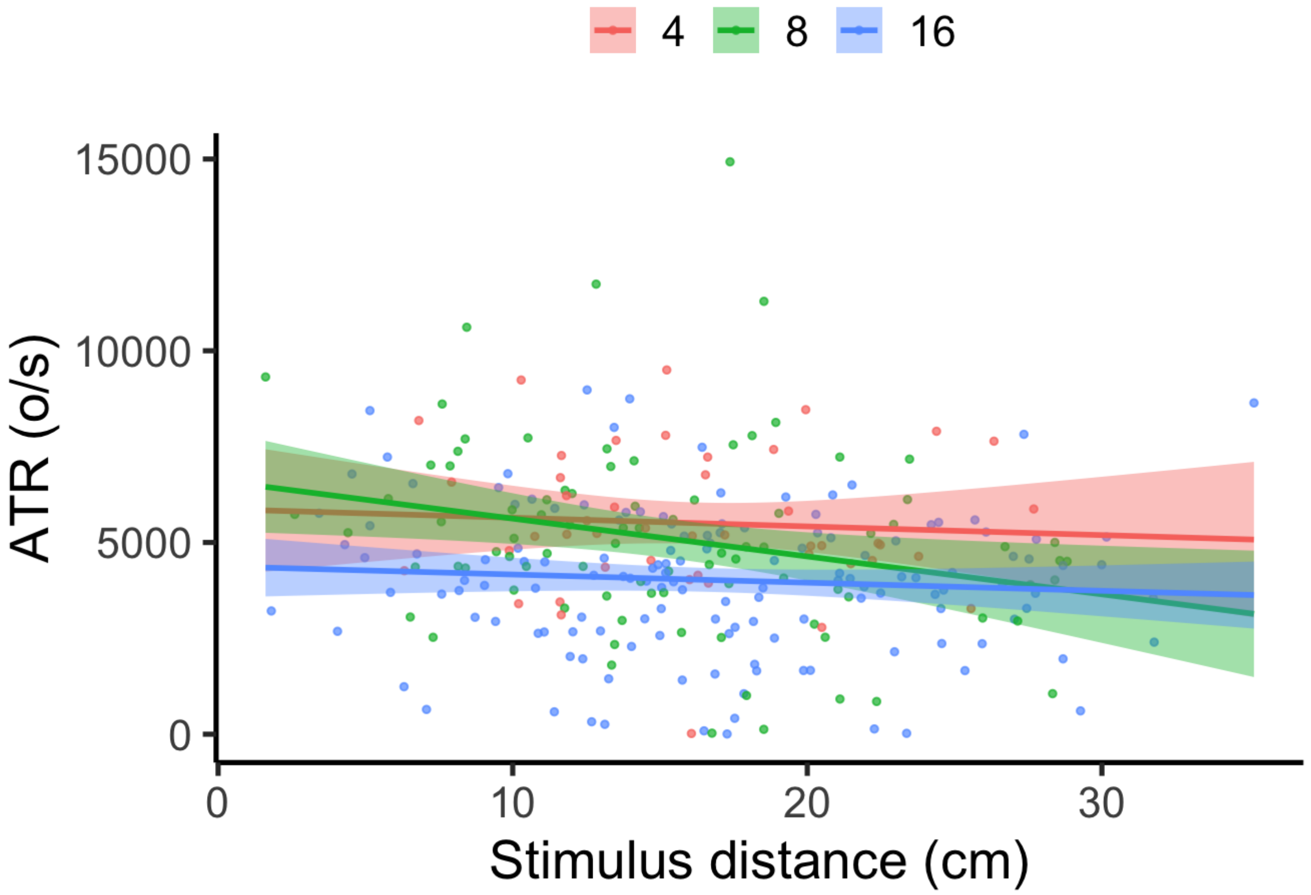


```
avt.emm <-emmeans(avt.lmer, ~Group.Size)
summary(avt.emm)
```

```
##  Group.Size emmean  SE   df lower.CL upper.CL
##  4            5476 394 52.9     4686     6266
##  8            4949 347 32.1     4242     5656
##  16           4068 321 23.6     3406     4730
##
## Degrees-of-freedom method: kenward-roger
## Confidence level used: 0.95
```

```
avt.emm<-as.data.frame(avt.emm)
avt.emplot = ggplot(avt.emm, aes(x=Group.Size, y=emmean, color=Group.Size, fill=Group.Si
ze)) +
  geom_pointrange(aes(ymin = lower.CL, ymax = upper.CL, color = Group.Size), na.rm = TRU
E, shape=21, size=.7, data = avt.emm) +
  geom_jitter(aes(x=Group.Size, y = Average.Turning.Rate.per.s, group=Group.Size), shape
= 21, size=1.2, width=.25, alpha = 0.35, data = fs_noNA) +
  xlab(bquote('Group Size')) + ylab(bquote('ATR (o/s)')) +
  theme_bw() + theme(panel.grid.major = element_blank(), panel.grid.minor = element_blan
k()) +
  theme(axis.text = element_text(size = 12)) + theme(axis.title = element_text(size = 1
2)) +
  theme_classic(base_family='Arial', base_size = 28) +
  theme(legend.position = "none")

avt.emplot
```

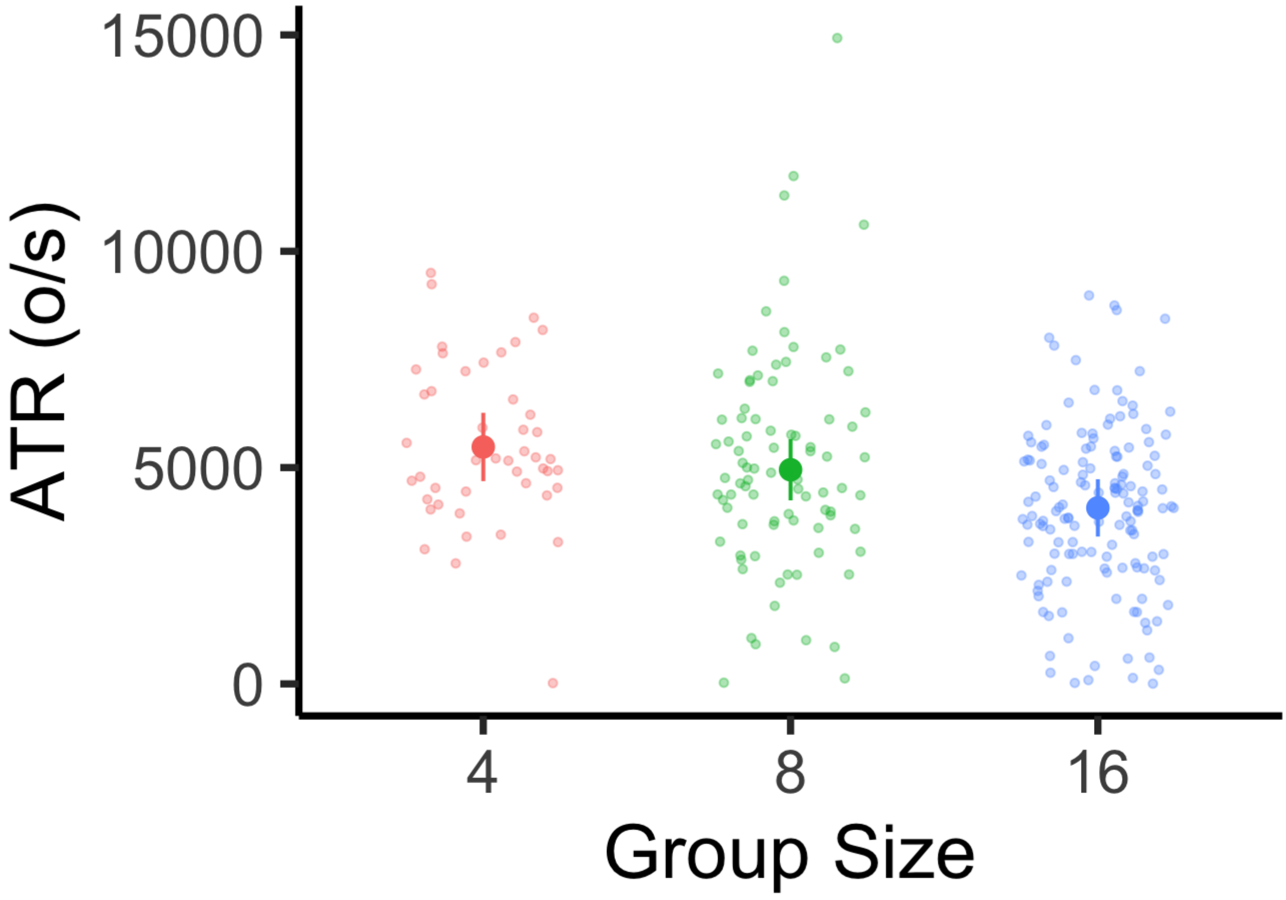


```
#DC
dc.lmer = lmer(Distance.covered ~ Group.Size*Stimulus.distance + (1|Video), data=fs_noN
A, na.action = "na.omit")
plot(dc.lmer)
```

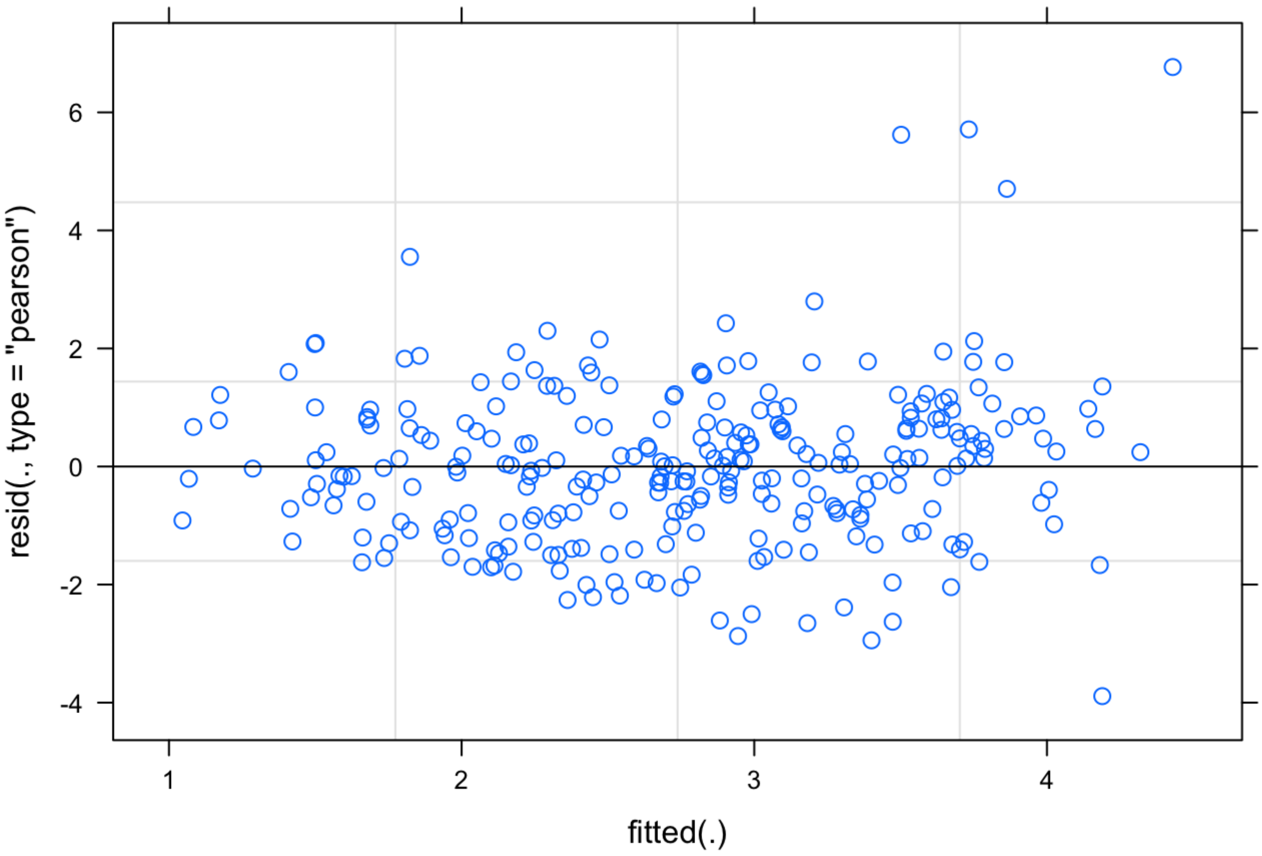


```
qqnorm(resid(dc.lmer)) #needs a boxcox transformation
qqline(resid(dc.lmer))
```

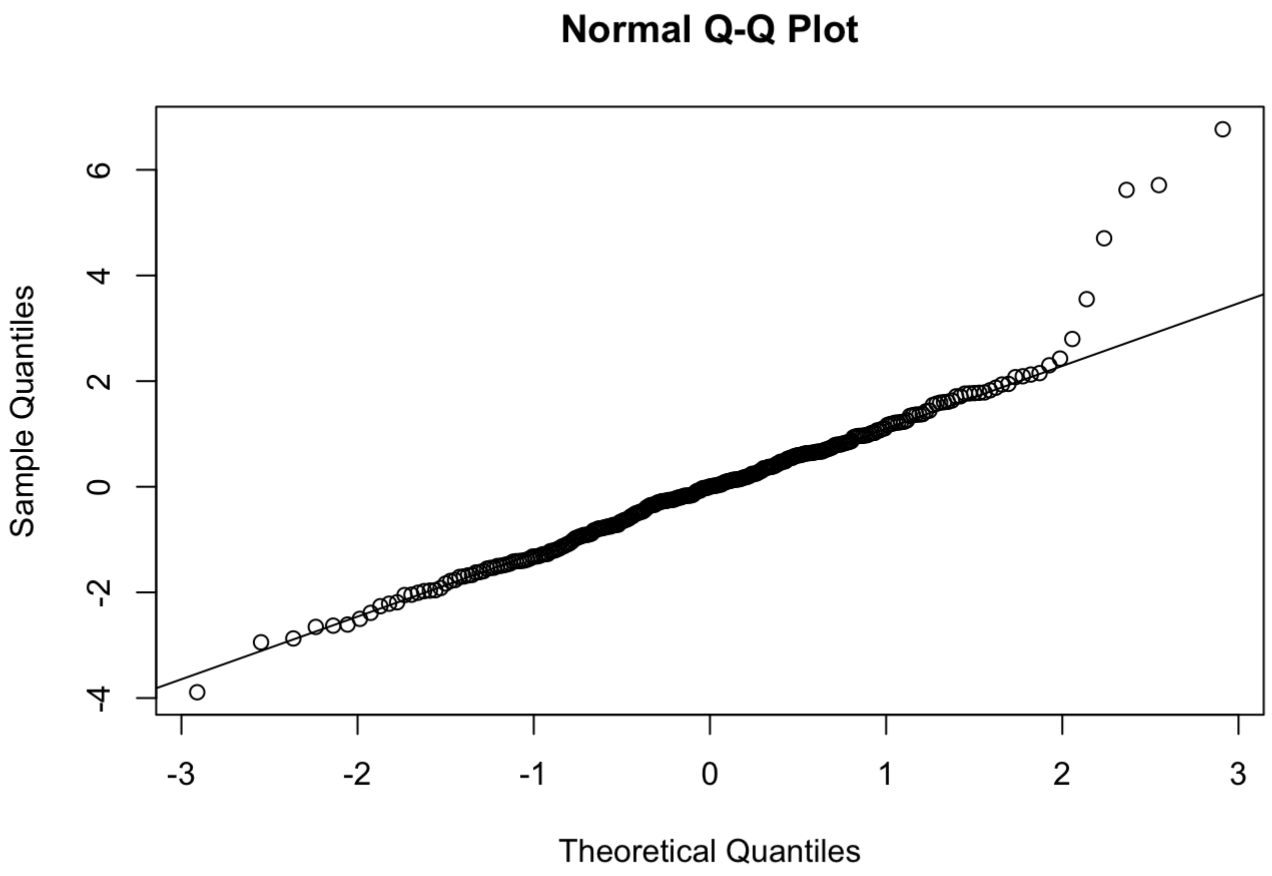


```
shapiro.test(resid(dc.lmer)) #p-value = 1.902e-08 so non normal
```

```
##
##  Shapiro-Wilk normality test
##
## data:  resid(dc.lmer)
## W = 0.94708, p-value = 1.902e-08
```

```
bartlett.test(resid(dc.lmer), fs_noNA$Group.Size) # p-value = 3.129e-05 so heteroscedast
ic
```

```
##
##  Bartlett test of homogeneity of variances
##
## data:  resid(dc.lmer) and fs_noNA$Group.Size
## Bartlett's K-squared = 20.744, df = 2, p-value = 3.129e-05
```

```
dc.lmer.bc <- powerTransform(dc.lmer, family="bcPower")
summary(dc.lmer.bc) #both significant
```

```
## bcPower Transformation to Normality
##       Est Power Rounded Pwr Wald Lwr Bnd Wald Upr Bnd
## [1,]    0.5941         0.5       0.4694       0.7188
##
## Likelihood ratio test that transformation parameter is equal to 0
##  (log transformation)
##                              LRT df       pval
## LR test, lambda = (0) 104.0107  1 < 2.22e-16
##
## Likelihood ratio test that no transformation is needed
##                              LRT df     pval
## LR test, lambda = (1) 37.33074  1 9.97e-10
```

```
dc.lmer.bc$roundlam
```

```
## [1] 0.5
```

```
dc.bc.lmer = lmer(bcPower(Distance.covered, dc.lmer.bc$roundlam) ~  Group.Size*Stimulus.
distance + (1|Video), data=fs_noNA, na.action = "na.omit")
plot(dc.bc.lmer)
```

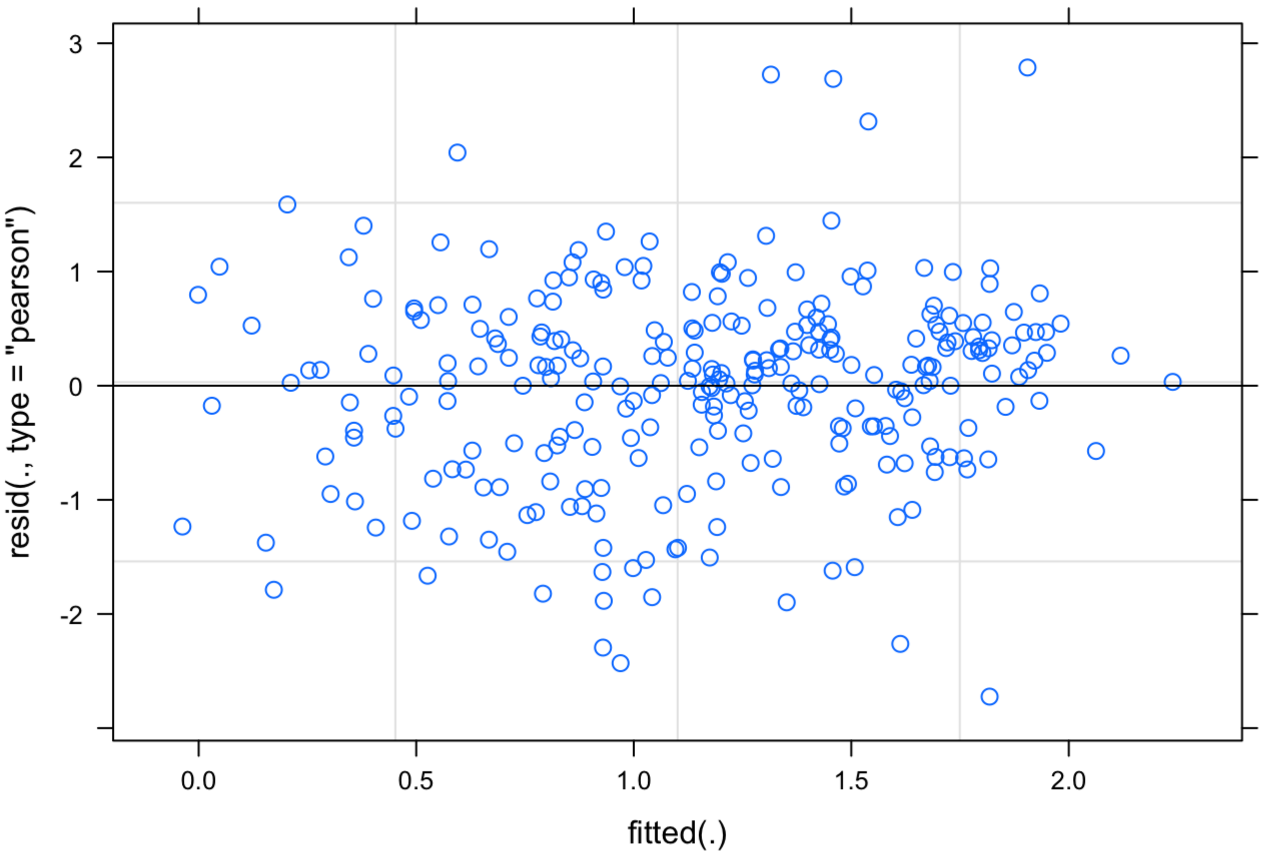

```
qqnorm(resid(dc.bc.lmer))
qqline(resid(dc.bc.lmer))
```

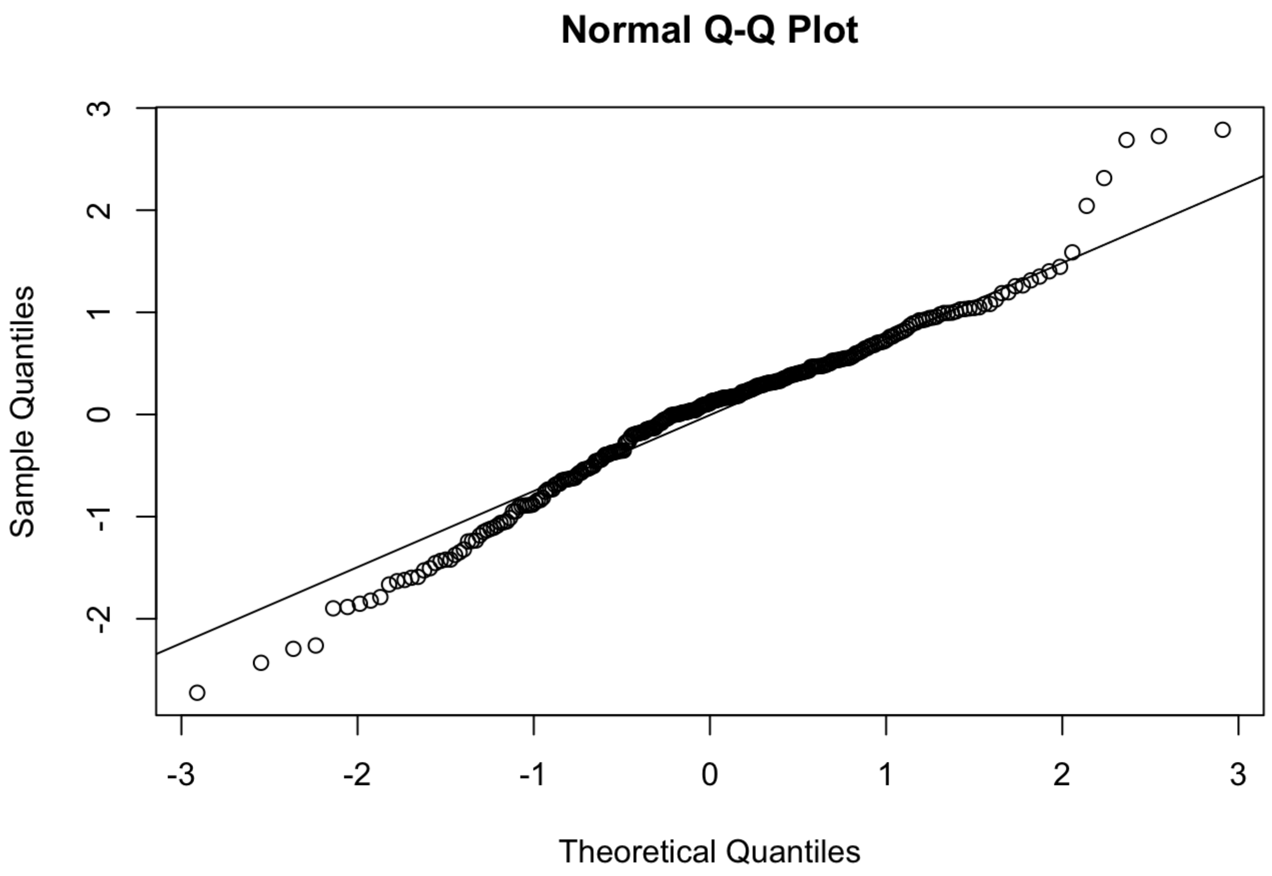


```
shapiro.test(resid(dc.bc.lmer))  #p=0.0001244 - glmer robust to minor deviations in assu
mptions so continuing with this transformed/improved model
```

```
##
##  Shapiro-Wilk normality test
##
## data:  resid(dc.bc.lmer)
## W = 0.97587, p-value = 0.0001244
```

```
bartlett.test(resid(dc.bc.lmer), fs_noNA$Group.Size) #p= 0.002 but visual inspection of
the plot looks fine so disregarding
```

```
##
##  Bartlett test of homogeneity of variances
##
## data:  resid(dc.bc.lmer) and fs_noNA$Group.Size
## Bartlett's K-squared = 12.796, df = 2, p-value = 0.001665
```

```
dc.bc.lmer2 = lmer(bcPower(Distance.covered, dc.lmer.bc$roundlam) ~  Group.Size+Stimulu
s.distance + (1|Video), data=fs_noNA, na.action = "na.omit")
anova(dc.bc.lmer, dc.bc.lmer2) #simpler lmer2 only has an AIC that is lower by <1 so sti
cking with most complex model
```

```
## Data: fs_noNA
## Models:
## dc.bc.lmer2: bcPower(Distance.covered, dc.lmer.bc$roundlam) ~ Group.Size + Stimulus.d
istance + (1 | Video)
## dc.bc.lmer: bcPower(Distance.covered, dc.lmer.bc$roundlam) ~ Group.Size * Stimulus.di
stance + (1 | Video)
##             npar    AIC    BIC  logLik deviance  Chisq Df Pr(>Chisq)
## dc.bc.lmer2    6 758.78 780.53 -373.39   746.78                     
## dc.bc.lmer     8 759.63 788.63 -371.82   743.63 3.1504  2      0.207
```

```
summary(dc.bc.lmer)
```

```
## Linear mixed model fit by REML ['lmerMod']
## Formula: bcPower(Distance.covered, dc.lmer.bc$roundlam) ~ Group.Size *
##     Stimulus.distance + (1 | Video)
##    Data: fs_noNA
##
## REML criterion at convergence: 768.8
##
## Scaled residuals:
##     Min      1Q  Median      3Q     Max
## -3.0578 -0.5689  0.1244  0.5591  3.1289
##
## Random effects:
##  Groups   Name        Variance Std.Dev.
##  Video    (Intercept) 0.1266   0.3559
##  Residual             0.7937   0.8909
## Number of obs: 277, groups:  Video, 36
##
## Fixed effects:
##                                Estimate Std. Error t value
## (Intercept)                     1.97831    0.47175   4.194
## Group.Size8                     0.35817    0.55206   0.649
## Group.Size16                   -0.18039    0.52360  -0.345
## Stimulus.distance              -0.01800    0.02721  -0.661
## Group.Size8:Stimulus.distance  -0.05385    0.03173  -1.697
## Group.Size16:Stimulus.distance -0.03175    0.02941  -1.080
##
## Correlation of Fixed Effects:
##             (Intr) Grp.S8 Gr.S16 Stmls. G.S8:S
## Group.Size8 -0.855
## Group.Siz16 -0.901  0.770
## Stmls.dstnc -0.935  0.799  0.843
## Grp.Sz8:St.  0.802 -0.918 -0.723 -0.857
## Grp.Sz16:S.  0.865 -0.739 -0.916 -0.925  0.793
```

```
Anova(dc.bc.lmer, test="F")
```

```
## Analysis of Deviance Table (Type II Wald F tests with Kenward-Roger df)
##
## Response: bcPower(Distance.covered, dc.lmer.bc$roundlam)
##                                   F Df  Df.res    Pr(>F)
## Group.Size                   5.5138  2  34.874  0.008304 **
## Stimulus.distance           36.0588  1 270.698 6.141e-09 ***
## Group.Size:Stimulus.distance  1.5097  2 263.398  0.222877
## ---
## Signif. codes:  0 '***' 0.001 '**' 0.01 '*' 0.05 '.' 0.1 ' ' 1
```

```
r.squaredGLMM(dc.bc.lmer)
```

```
##            R2m       R2c
## [1,] 0.1789889 0.2919535
```

```
backtrans.dc <- update(ref_grid(dc.bc.lmer), tran = make.tran("boxcox", 0.5)) #back tran
sforms from boxcox so emmeans produces sensible estimates of means and SE where 0.5 is t
he lambda estimated by powertransform function above
emmeans(backtrans.dc, pairwise~Group.Size, adjust = c("Tukey"), type = "response")
```

```
## $emmeans
##  Group.Size response    SE   df lower.CL upper.CL
##  4              3.40 0.308 62.5     2.81     4.04
##  8              2.51 0.224 32.7     2.07     2.98
##  16             2.23 0.190 21.5     1.85     2.64
##
## Degrees-of-freedom method: kenward-roger
## Confidence level used: 0.95
## Intervals are back-transformed from the Box-Cox (lambda = 0.5) scale
##
## $contrasts
##  contrast                  estimate    SE   df t.ratio p.value
##  Group.Size4 - Group.Size8     0.519 0.219 46.7   2.368  0.0564
##  Group.Size4 - Group.Size16    0.697 0.210 39.9   3.323  0.0053
##  Group.Size8 - Group.Size16    0.179 0.190 26.9   0.939  0.6208
##
## Note: contrasts are still on the Box-Cox (lambda = 0.5) scale
## Degrees-of-freedom method: kenward-roger
## P value adjustment: tukey method for comparing a family of 3 estimates
```

```
dc.emm<-emmeans(backtrans.dc, ~Group.Size, type = "response")
summary(dc.emm)
```

```
##  Group.Size response    SE   df lower.CL upper.CL
##  4              3.40 0.308 62.5     2.81     4.04
##  8              2.51 0.224 32.7     2.07     2.98
##  16             2.23 0.190 21.5     1.85     2.64
##
## Degrees-of-freedom method: kenward-roger
## Confidence level used: 0.95
## Intervals are back-transformed from the Box-Cox (lambda = 0.5) scale
```

```
dc.emm<-as.data.frame(dc.emm)
dc.emplot = ggplot(dc.emm, aes(x=Group.Size, y=response, color=Group.Size, fill=Group.Si
ze)) +
  geom_pointrange(aes(ymin = lower.CL, ymax = upper.CL, color = Group.Size), na.rm = TRU
E, shape=21, size=.7, data = dc.emm) +
  geom_jitter(aes(x=Group.Size, y = Distance.covered, group=Group.Size), shape = 21, siz
e=1.2, width=.25, alpha = 0.35, data = fs_noNA) +
  xlab(bquote('Group Size')) + ylab(bquote('Distance covered (mm)')) +
  theme_bw() + theme(panel.grid.major = element_blank(), panel.grid.minor = element_blan
k()) +
  theme(axis.text = element_text(size = 12)) + theme(axis.title = element_text(size = 1
2)) +
  theme_classic(base_family='Arial', base_size = 28) +
  theme(legend.position = "none")

dc.emplot
```

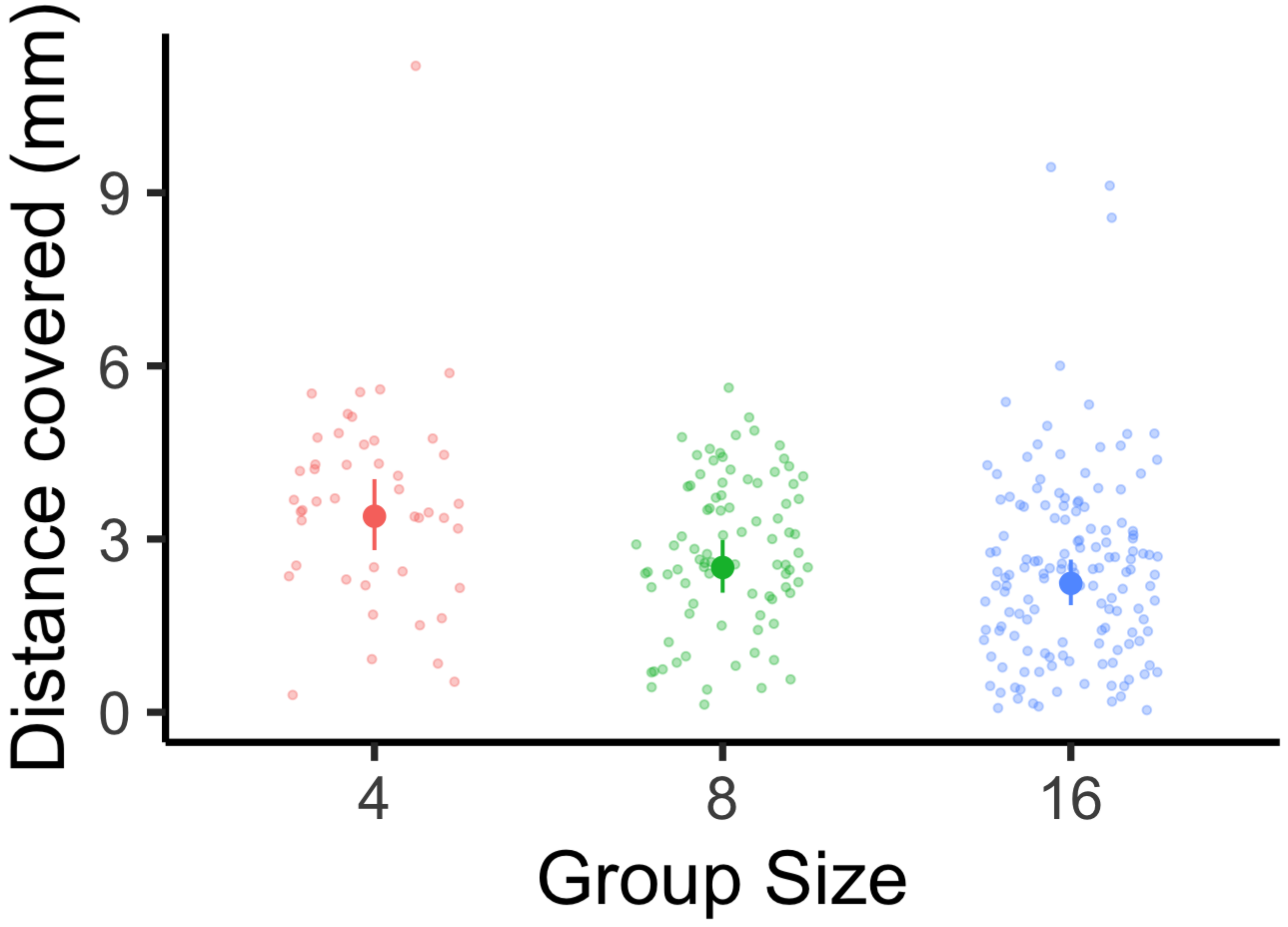

```
dc.plot <- ggplot(fs_noNA, aes(x=Stimulus.distance, y=Distance.covered, colour=Group.Siz
e)) +
  geom_point(aes(color = Group.Size), alpha = 0.7, shape=21, size=.7) +
  geom_smooth(method="glm", fullrange=TRUE, aes(fill = Group.Size)) +
  xlab(bquote('Stimulus distance (cm)')) + ylab(bquote('Distance covered (mm)')) +
  theme_classic(base_family='Arial', base_size = 28) +
  theme(legend.title=element_blank()) +
  theme(legend.position = "top")
dc.plot
```

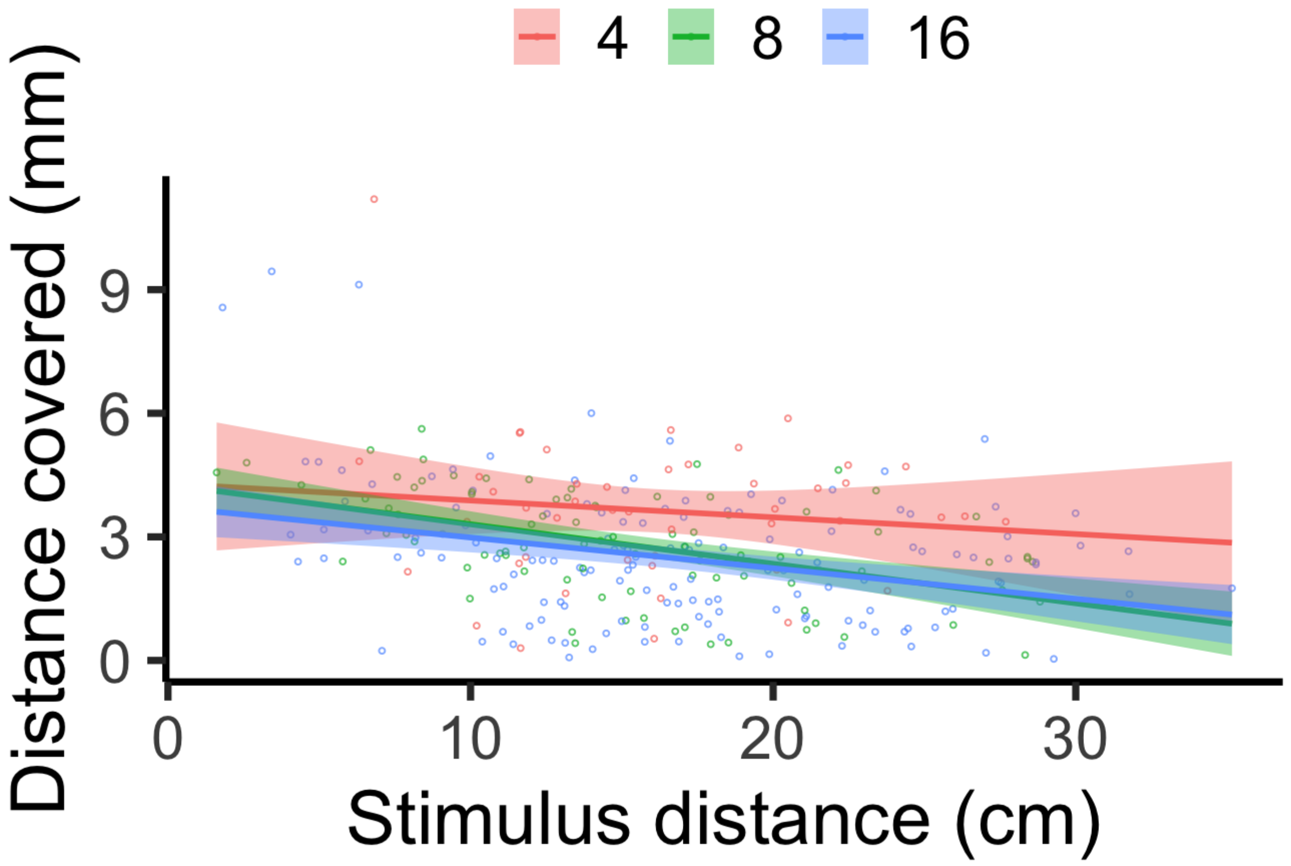


### SCHOOL BEHAVIOR AFTER STIMULATION

Throughout the response, the school's cohesion and coordination were measured through nearest neighbor distance and alignment, respectively. Nearest neighbor distance (NND) represents the distance between each fish's center of mass and their most proximal neighbor's center of mass in the school. Alignment measures the variation in each school's members orientation with respect to the water's flow (0°). This variation was quantified by using the program Oriana 4 and determining each school members' angle then calculating the length of mean circular vector (r) of the group, which ranges from 0 (all members are at random angles) to 1 (all angles coordinated exactly). These characteristics were examined at intervals following the stimulus, including 0 ms (representing the school's cohesiveness and coordination immediately prior to the stimulus), 30 ms (representing the typical time for this species to complete stage 1), and 100 ms (the average time for individuals to complete both stages 1 and 2).

```
#NND
NND1 <- read.csv("/Users/laurennadler/Desktop/MyRData/Monica_groupsize_NND.csv", header=
T)
NND1$Groupsize = as.factor(NND1$Groupsize)
NND1$Time = as.factor(NND1$Time)
nnd.lmer = lmer(NND ~  Groupsize*Time + (1|Video/Fish), data=NND1, na.action = "na.omi
t")
summary(nnd.lmer)
```

```
## Linear mixed model fit by REML ['lmerMod']
## Formula: NND ~ Groupsize * Time + (1 | Video/Fish)
##    Data: NND1
## 
## REML criterion at convergence: 4536.7
## 
## Scaled residuals: 
##     Min      1Q  Median      3Q     Max 
## -3.5519 -0.4871 -0.0523  0.4095  6.1185 
## 
## Random effects:
##  Groups     Name        Variance Std.Dev.
##  Fish:Video (Intercept) 3.8462   1.9612  
##  Video      (Intercept) 0.1788   0.4229  
##  Residual               3.1169   1.7655  
## Number of obs: 1008, groups:  Fish:Video, 336; Video, 36
## 
## Fixed effects:
##                     Estimate Std. Error t value
## (Intercept)           6.2305     0.4000  15.578
## Groupsize8           -1.1012     0.4974  -2.214
## Groupsize16          -1.8928     0.4595  -4.119
## Time30                0.1556     0.3604   0.432
## Time100               1.9922     0.3604   5.528
## Groupsize8:Time30    -0.2278     0.4414  -0.516
## Groupsize16:Time30   -0.1681     0.4029  -0.417
## Groupsize8:Time100   -1.7111     0.4414  -3.877
## Groupsize16:Time100  -1.4165     0.4029  -3.516
## 
## Correlation of Fixed Effects:
##             (Intr) Grpsz8 Grps16 Time30 Tim100 G8:T30 G16:T3 G8:T10
## Groupsize8  -0.804                                                 
## Groupsize16 -0.870  0.700                                          
## Time30      -0.451  0.362  0.392                                   
## Time100     -0.451  0.362  0.392  0.500                            
## Grpsz8:Tm30  0.368 -0.444 -0.320 -0.816 -0.408                     
## Grpsz16:T30  0.403 -0.324 -0.438 -0.894 -0.447  0.730              
## Grpsz8:T100  0.368 -0.444 -0.320 -0.408 -0.816  0.500  0.365       
## Grps16:T100  0.403 -0.324 -0.438 -0.447 -0.894  0.365  0.500  0.730
```

```
plot(nnd.lmer) #funneled
```

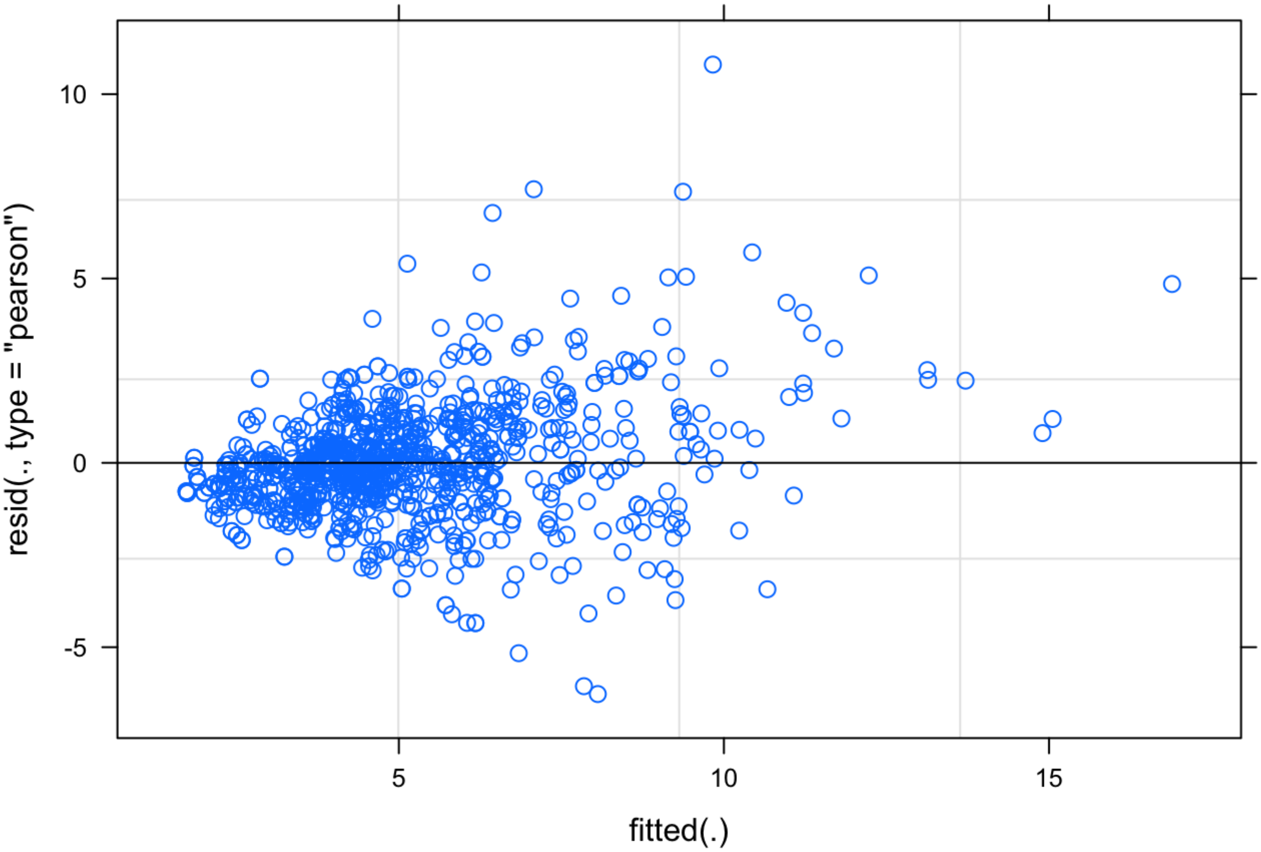


```
qqnorm(resid(nnd.lmer)) #curvy
qqline(resid(nnd.lmer))
```

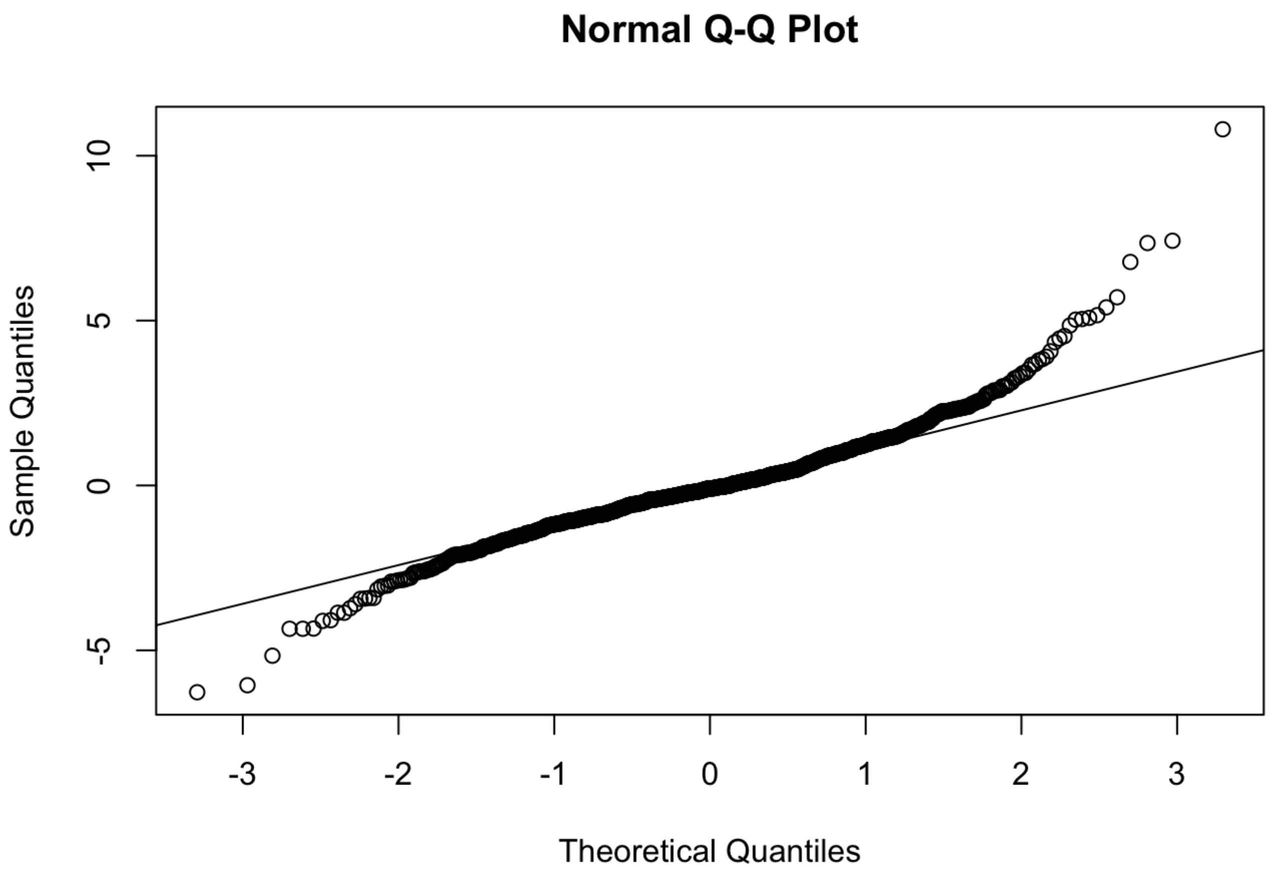


```
shapiro.test(resid(nnd.lmer)) #p < 0.001
```

```
##
##  Shapiro-Wilk normality test
##
## data:  resid(nnd.lmer)
## W = 0.94188, p-value < 2.2e-16
```

```
bartlett.test(resid(nnd.lmer), NND1$Groupsize) #p-value < 0.001
```

```
##
##  Bartlett test of homogeneity of variances
##
## data:  resid(nnd.lmer) and NND1$Groupsize
## Bartlett's K-squared = 61.391, df = 2, p-value = 4.668e-14
```

```
nnd.lmer.bc <- powerTransform(nnd.lmer, family="bcPower")
summary(nnd.lmer.bc) #both significant
```

```
## bcPower Transformation to Normality
##      Est Power Rounded Pwr Wald Lwr Bnd Wald Upr Bnd
## [1,]    0.2964        0.33        0.215       0.3777
##
## Likelihood ratio test that transformation parameter is equal to 0
##  (log transformation)
##                            LRT df       pval
## LR test, lambda = (0) 52.75842  1 3.7725e-13
##
## Likelihood ratio test that no transformation is needed
##                            LRT df       pval
## LR test, lambda = (1) 268.0465  1 < 2.22e-16
```

```
nnd.lmer.bc$roundlam
```

```
## [1] 0.33
```

```
nnd.bc.lmer = lmer(bcPower(NND, nnd.lmer.bc$roundlam) ~  Groupsize*Time + (1|Video/Fis
h), data=NND1, na.action = "na.omit")
plot(nnd.bc.lmer) #much better
```

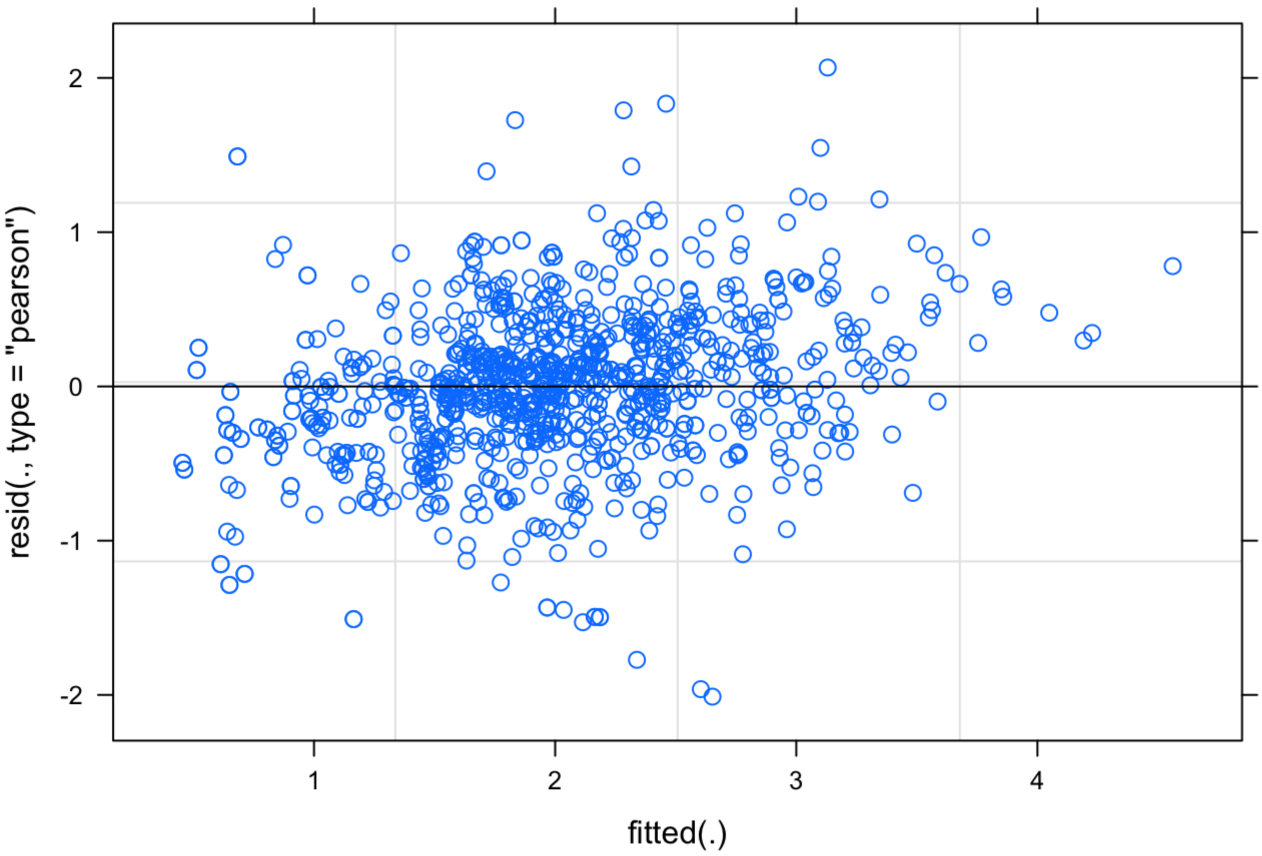

```
qqnorm(resid(nnd.bc.lmer)) #much improved
qqline(resid(nnd.bc.lmer))
```

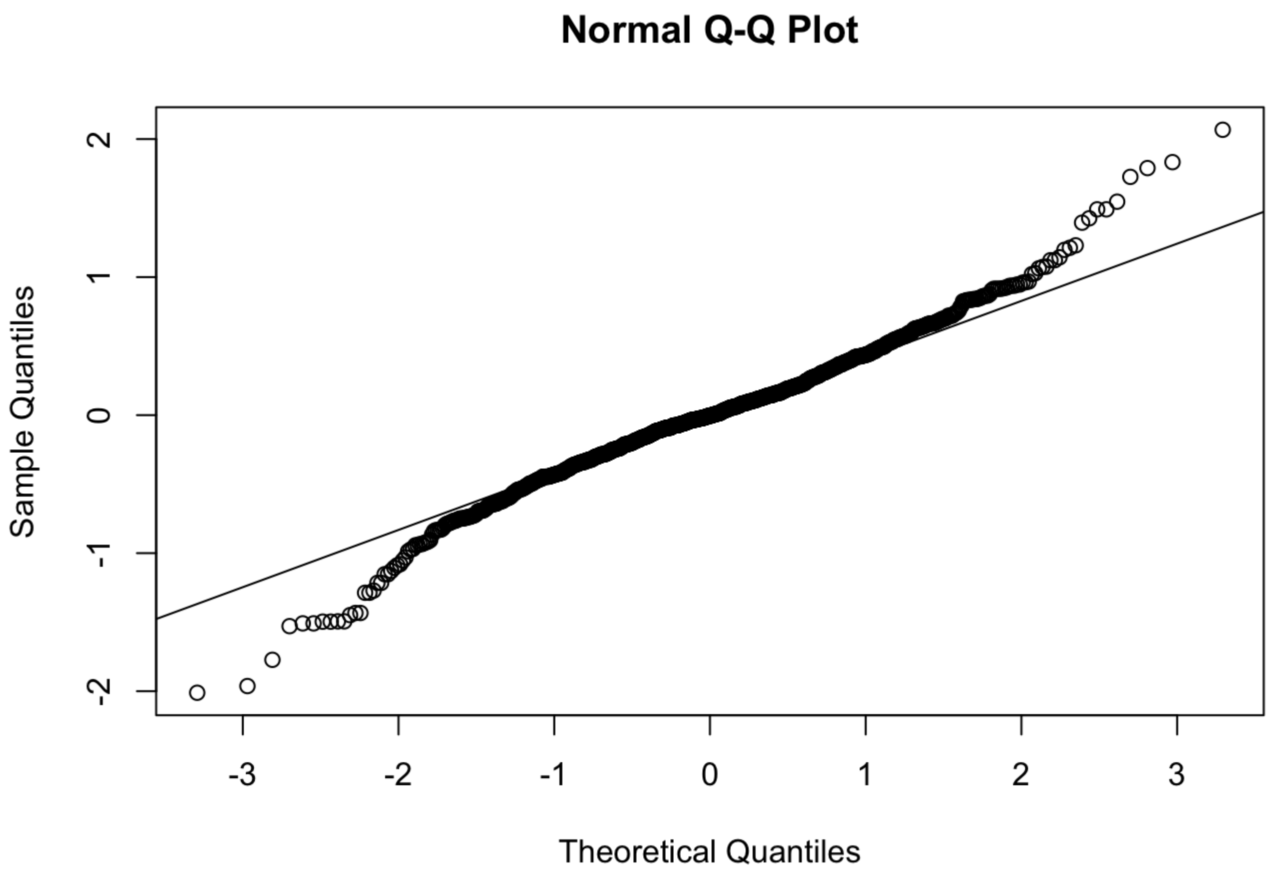


```
shapiro.test(resid(nnd.bc.lmer)) #p-value = 5.897e-10, but qqplot much improved so conti
nuing with this model
```

```
##
##  Shapiro-Wilk normality test
##
## data:  resid(nnd.bc.lmer)
## W = 0.98166, p-value = 5.897e-10
```

```
bartlett.test(resid(nnd.bc.lmer), NND1$Groupsize) #p-value = 3.469e-05, but residuals pl
ot much improved so continuing with this model
```

```
##
##  Bartlett test of homogeneity of variances
##
## data:  resid(nnd.bc.lmer) and NND1$Groupsize
## Bartlett's K-squared = 20.538, df = 2, p-value = 3.469e-05
```

```
bartlett.test(resid(nnd.bc.lmer), NND1$Time) #p-value < 2.2e-16, but residuals plot much improved so continuing with this model
```

```
##
##  Bartlett test of homogeneity of variances
##
## data:  resid(nnd.bc.lmer) and NND1$Time
## Bartlett's K-squared = 153.77, df = 2, p-value < 2.2e-16
```

```
nnd.bc.lmer2 = lmer(bcPower(NND, nnd.lmer.bc$roundlam) ~  Groupsize+Time + (1|Video/Fis
h), data=NND1, na.action = "na.omit")
anova(nnd.bc.lmer, nnd.bc.lmer2) #simpler model lmer2 lower AIC but <2 so keep more comp
lex model
```

```
## Data: NND1
## Models:
## nnd.bc.lmer2: bcPower(NND, nnd.lmer.bc$roundlam) ~ Groupsize + Time + (1 | Video/Fis
h)
## nnd.bc.lmer: bcPower(NND, nnd.lmer.bc$roundlam) ~ Groupsize * Time + (1 | Video/Fish)
##              npar    AIC    BIC  logLik deviance  Chisq Df Pr(>Chisq)
## nnd.bc.lmer2    8 2291.2 2330.5 -1137.6   2275.2
## nnd.bc.lmer    12 2292.8 2351.8 -1134.4   2268.8 6.4154  4     0.1702
```

```
summary(nnd.bc.lmer)
```

```
## Linear mixed model fit by REML ['lmerMod']
## Formula: bcPower(NND, nnd.lmer.bc$roundlam) ~ Groupsize * Time + (1 |
##     Video/Fish)
##    Data: NND1
##
## REML criterion at convergence: 2297.5
##
## Scaled residuals:
##     Min      1Q  Median      3Q     Max
## -3.4971 -0.4901 -0.0069  0.4829  3.5949
##
## Random effects:
##  Groups     Name        Variance Std.Dev.
##  Fish:Video (Intercept) 0.4026   0.6345
##  Video      (Intercept) 0.0328   0.1811
##  Residual               0.3307   0.5751
## Number of obs: 1008, groups:  Fish:Video, 336; Video, 36
##
## Fixed effects:
##                     Estimate Std. Error t value
## (Intercept)          2.40974    0.13421  17.956
## Groupsize8          -0.36670    0.16847  -2.177
## Groupsize16         -0.66054    0.15673  -4.215
## Time30               0.03542    0.11739   0.302
## Time100              0.36965    0.11739   3.149
## Groupsize8:Time30   -0.07197    0.14377  -0.501
## Groupsize16:Time30  -0.04302    0.13124  -0.328
## Groupsize8:Time100  -0.33645    0.14377  -2.340
## Groupsize16:Time100 -0.17848    0.13124  -1.360
##
## Correlation of Fixed Effects:
##             (Intr) Grpsz8 Grps16 Time30 Tim100 G8:T30 G16:T3 G8:T10
## Groupsize8  -0.797
## Groupsize16 -0.856  0.682
## Time30      -0.437  0.348  0.374
## Time100     -0.437  0.348  0.374  0.500
## Grpsz8:Tm30  0.357 -0.427 -0.306 -0.816 -0.408
## Grpsz16:T30  0.391 -0.312 -0.419 -0.894 -0.447  0.730
## Grpsz8:T100  0.357 -0.427 -0.306 -0.408 -0.816  0.500  0.365
## Grps16:T100  0.391 -0.312 -0.419 -0.447 -0.894  0.365  0.500  0.730
```

```
Anova(nnd.bc.lmer, test="F")
```

```
## Analysis of Deviance Table (Type II Wald F tests with Kenward-Roger df)
##
## Response: bcPower(NND, nnd.lmer.bc$roundlam)
##                      F Df Df.res    Pr(>F)
## Groupsize      14.3187  2  40.47 1.984e-05 ***
## Time           10.5609  2 666.00 3.053e-05 ***
## Groupsize:Time  1.5971  4 666.00    0.1733
## ---
## Signif. codes:  0 '***' 0.001 '**' 0.01 '*' 0.05 '.' 0.1 ' ' 1
```

```
r.squaredGLMM(nnd.bc.lmer)
```

```
##            R2m       R2c
## [1,] 0.0861159 0.6055021
```

```
backtrans.nnd <- update(ref_grid(nnd.bc.lmer), tran = make.tran("boxcox", 0.33)) #where
0.33 is the lambda estimated by powertransform function above
emmeans(backtrans.nnd, pairwise~Groupsize, adjust = c("Tukey"), type = "response")
```

```
## $emmeans
##  Groupsize response    SE    df lower.CL upper.CL
##  4             6.34 0.399 100.8     5.58     7.17
##  8             4.76 0.256  39.5     4.26     5.30
##  16            4.13 0.190  17.6     3.75     4.55
##
## Results are averaged over the levels of: Time
## Degrees-of-freedom method: kenward-roger
## Confidence level used: 0.95
## Intervals are back-transformed from the Box-Cox (lambda = 0.33) scale
##
## $contrasts
##  contrast                 estimate    SE   df t.ratio p.value
##  Groupsize4 - Groupsize8     0.503 0.147 68.1   3.430  0.0029
##  Groupsize4 - Groupsize16    0.734 0.137 53.1   5.353  <.0001
##  Groupsize8 - Groupsize16    0.232 0.116 27.5   1.994  0.1326
##
## Results are averaged over the levels of: Time
## Note: contrasts are still on the Box-Cox (lambda = 0.33) scale
## Degrees-of-freedom method: kenward-roger
## P value adjustment: tukey method for comparing a family of 3 estimates
```

```
emmeans(backtrans.nnd, pairwise~Groupsize*Time, adjust = c("Tukey"), type = "response")
```

```
## $emmeans
##  Groupsize Time response    SE    df lower.CL upper.CL
##  4         0        5.89 0.440 178.4     5.06     6.80
##  8         0        4.77 0.290  64.8     4.21     5.37
##  16        0        3.98 0.204  25.9     3.57     4.41
##  4         30       6.01 0.446 178.4     5.17     6.93
##  8         30       4.66 0.286  64.8     4.12     5.26
##  16        30       3.96 0.204  25.9     3.56     4.39
##  4         100      7.19 0.503 178.4     6.24     8.23
##  8         100      4.86 0.294  64.8     4.30     5.47
##  16        100      4.48 0.221  25.9     4.04     4.95
##
## Degrees-of-freedom method: kenward-roger
## Confidence level used: 0.95
## Intervals are back-transformed from the Box-Cox (lambda = 0.33) scale
##
## $contrasts
##  contrast                                  estimate     SE    df t.ratio p.value
##  Groupsize4 Time0 - Groupsize8 Time0         0.3667 0.1685 117.5   2.177  0.4277
##  Groupsize4 Time0 - Groupsize16 Time0        0.6605 0.1567  89.7   4.215  0.0019
##  Groupsize4 Time0 - Groupsize4 Time30       -0.0354 0.1174 666.0  -0.302  1.0000
##  Groupsize4 Time0 - Groupsize8 Time30        0.4032 0.1685 117.5   2.394  0.2974
##  Groupsize4 Time0 - Groupsize16 Time30       0.6681 0.1567  89.7   4.263  0.0016
##  Groupsize4 Time0 - Groupsize4 Time100      -0.3696 0.1174 666.0  -3.149  0.0448
##  Groupsize4 Time0 - Groupsize8 Time100       0.3335 0.1685 117.5   1.980  0.5606
##  Groupsize4 Time0 - Groupsize16 Time100      0.4694 0.1567  89.7   2.995  0.0811
##  Groupsize8 Time0 - Groupsize16 Time0        0.2938 0.1301  43.3   2.259  0.3885
##  Groupsize8 Time0 - Groupsize4 Time30       -0.4021 0.1685 117.5  -2.387  0.3010
##  Groupsize8 Time0 - Groupsize8 Time30        0.0366 0.0830 666.0   0.440  1.0000
##  Groupsize8 Time0 - Groupsize16 Time30       0.3014 0.1301  43.3   2.317  0.3548
##  Groupsize8 Time0 - Groupsize4 Time100      -0.7363 0.1685 117.5  -4.371  0.0009
##  Groupsize8 Time0 - Groupsize8 Time100      -0.0332 0.0830 666.0  -0.400  1.0000
##  Groupsize8 Time0 - Groupsize16 Time100      0.1027 0.1301  43.3   0.789  0.9966
##  Groupsize16 Time0 - Groupsize4 Time30      -0.6960 0.1567  89.7  -4.441  0.0008
##  Groupsize16 Time0 - Groupsize8 Time30      -0.2573 0.1301  43.3  -1.978  0.5663
##  Groupsize16 Time0 - Groupsize16 Time30      0.0076 0.0587 666.0   0.129  1.0000
##  Groupsize16 Time0 - Groupsize4 Time100     -1.0302 0.1567  89.7  -6.573  <.0001
##  Groupsize16 Time0 - Groupsize8 Time100     -0.3270 0.1301  43.3  -2.514  0.2539
##  Groupsize16 Time0 - Groupsize16 Time100    -0.1912 0.0587 666.0  -3.257  0.0322
##  Groupsize4 Time30 - Groupsize8 Time30       0.4387 0.1685 117.5   2.604  0.1964
##  Groupsize4 Time30 - Groupsize16 Time30      0.7036 0.1567  89.7   4.489  0.0007
##  Groupsize4 Time30 - Groupsize4 Time100     -0.3342 0.1174 666.0  -2.847  0.1039
##  Groupsize4 Time30 - Groupsize8 Time100      0.3689 0.1685 117.5   2.190  0.4192
##  Groupsize4 Time30 - Groupsize16 Time100     0.5048 0.1567  89.7   3.221  0.0446
##  Groupsize8 Time30 - Groupsize16 Time30      0.2649 0.1301  43.3   2.036  0.5280
##  Groupsize8 Time30 - Groupsize4 Time100     -0.7729 0.1685 117.5  -4.588  0.0004
##  Groupsize8 Time30 - Groupsize8 Time100     -0.0698 0.0830 666.0  -0.840  0.9956
##  Groupsize8 Time30 - Groupsize16 Time100     0.0661 0.1301  43.3   0.508  0.9999
##  Groupsize16 Time30 - Groupsize4 Time100    -1.0378 0.1567  89.7  -6.622  <.0001
##  Groupsize16 Time30 - Groupsize8 Time100    -0.3346 0.1301  43.3  -2.572  0.2280
##  Groupsize16 Time30 - Groupsize16 Time100   -0.1988 0.0587 666.0  -3.386  0.0213
##  Groupsize4 Time100 - Groupsize8 Time100     0.7031 0.1685 117.5   4.174  0.0018
```

```
##  Groupsize4 Time100 - Groupsize16 Time100   0.8390 0.1567  89.7   5.353  <.0001
##  Groupsize8 Time100 - Groupsize16 Time100   0.1359 0.1301  43.3   1.044  0.9789
##
## Note: contrasts are still on the Box-Cox (lambda = 0.33) scale
## Degrees-of-freedom method: kenward-roger
## P value adjustment: tukey method for comparing a family of 9 estimates
```

```
nnd.emm<-emmeans(backtrans.nnd, ~Groupsize*Time, type = "response")
summary(nnd.emm)
```

```
##  Groupsize Time response    SE    df lower.CL upper.CL
##  4         0        5.89 0.440 178.4     5.06     6.80
##  8         0        4.77 0.290  64.8     4.21     5.37
##  16        0        3.98 0.204  25.9     3.57     4.41
##  4         30       6.01 0.446 178.4     5.17     6.93
##  8         30       4.66 0.286  64.8     4.12     5.26
##  16        30       3.96 0.204  25.9     3.56     4.39
##  4         100      7.19 0.503 178.4     6.24     8.23
##  8         100      4.86 0.294  64.8     4.30     5.47
##  16        100      4.48 0.221  25.9     4.04     4.95
##
## Degrees-of-freedom method: kenward-roger
## Confidence level used: 0.95
## Intervals are back-transformed from the Box-Cox (lambda = 0.33) scale
```

```
nnd.emm<-as.data.frame(nnd.emm)
nnd.emplot = ggplot(nnd.emm, aes(x=Time, y=response, fill=Groupsize)) +
  geom_jitter(aes(x=Time, y = NND, colour=Groupsize), shape = 21, size=1, width=.35, alp
ha = 0.15, data = NND1) +
  geom_pointrange(aes(ymin = lower.CL, ymax = upper.CL, color = Groupsize), shape=21, si
ze=.2, data = nnd.emm) +
  geom_line(aes(group = Groupsize, colour = Groupsize), data = nnd.emm) +
  xlab(bquote('Time post-stimulus (ms)')) + ylab(bquote('NND (cm)')) +
  theme_bw() + theme(panel.grid.major = element_blank(), panel.grid.minor = element_blan
k()) +
  theme(axis.text = element_text(size = 12)) + theme(axis.title = element_text(size = 1
2)) +
  theme_classic(base_family='Arial', base_size = 28) +
  theme(legend.position = "none") +
  scale_y_log10()

nnd.emplot + facet_grid(. ~ Groupsize)
```

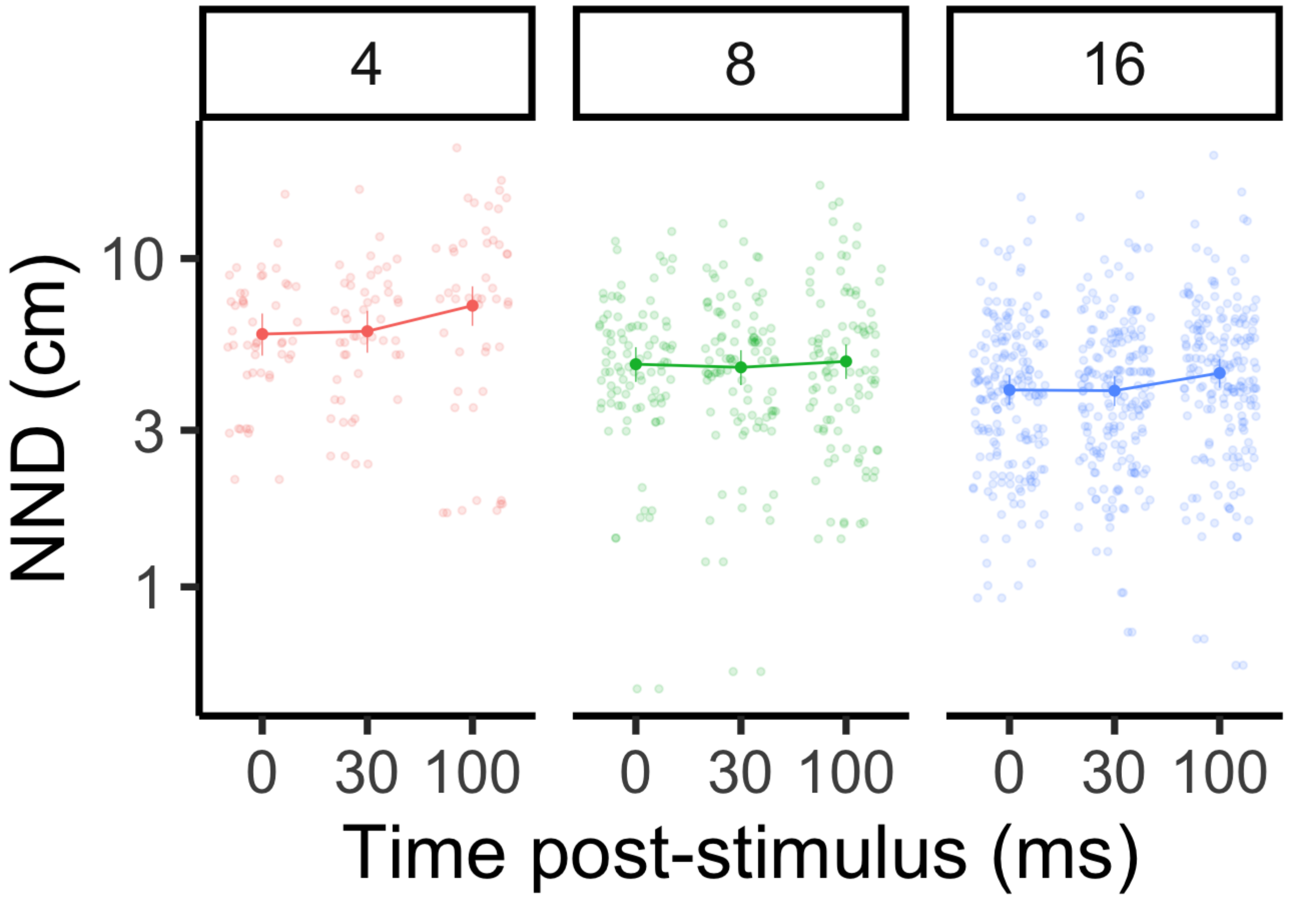


```
#Alignment
align <- read.csv("/Users/laurennadler/Desktop/MyRData/Monica_groupsize_alignment.csv",
header=T)
align$Groupsize = as.factor(align$Groupsize)
align$Time = as.factor(align$Time)

alignment.lmer = lmer(Alignment ~ Groupsize*Time + (1|Video), data=align, na.action = "n
a.omit")
plot(alignment.lmer)
```

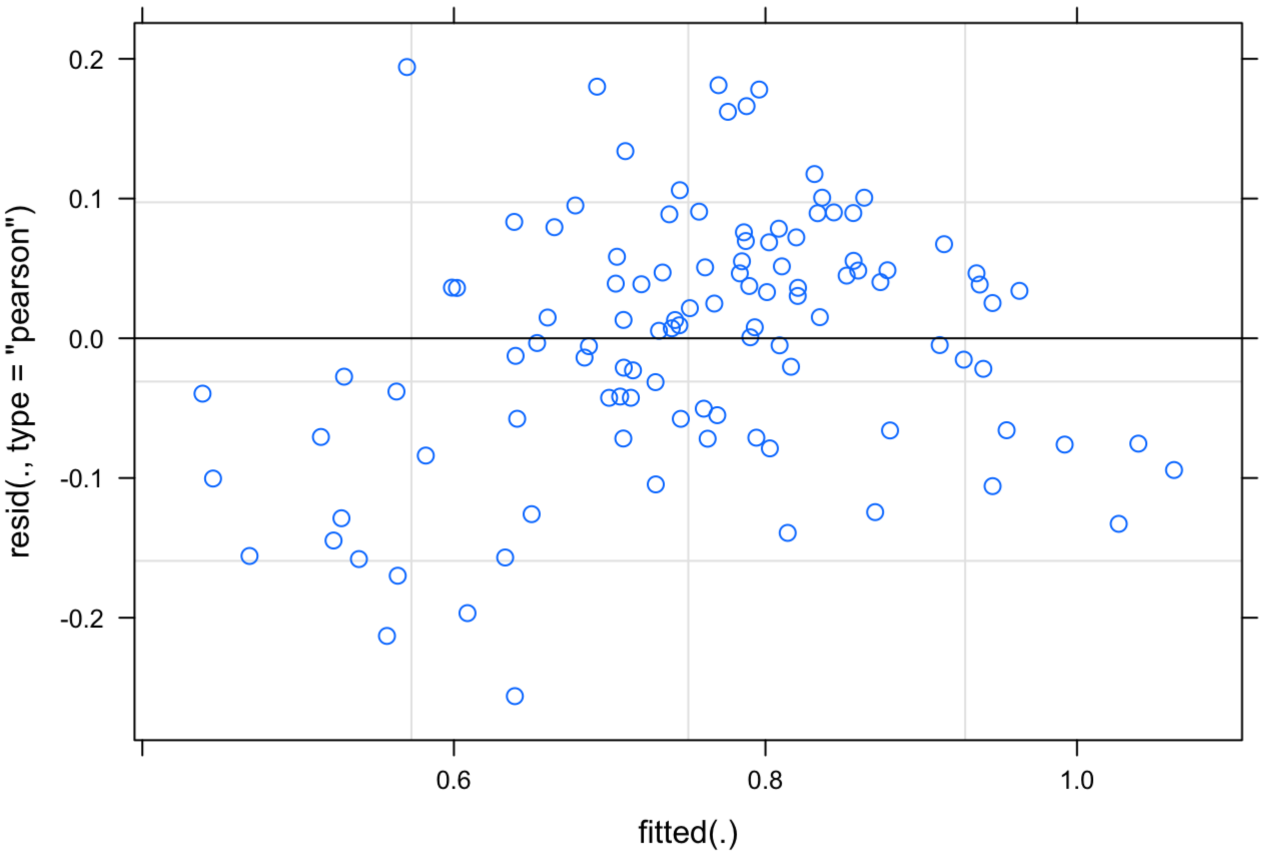


```
qqnorm(resid(alignment.lmer))
qqline(resid(alignment.lmer))
```

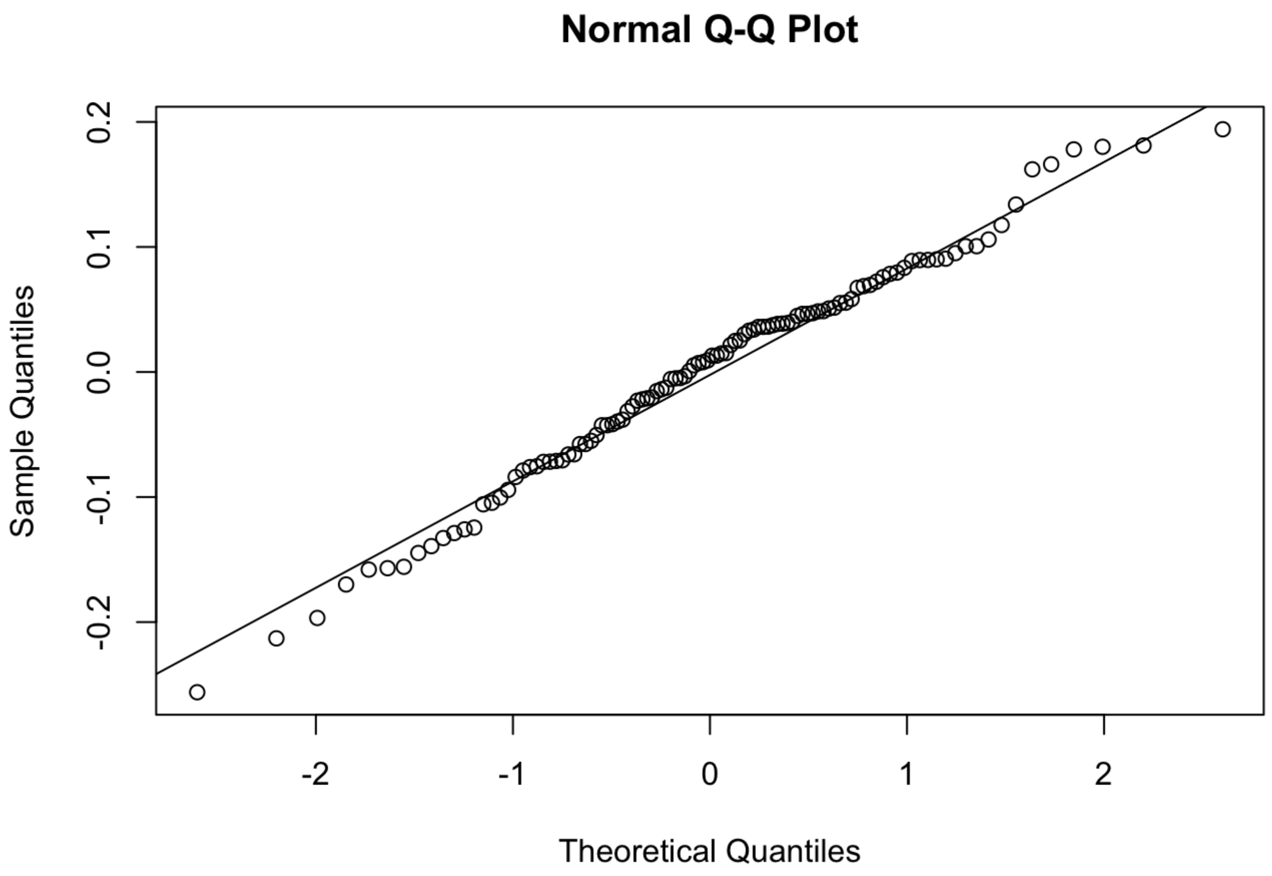


```
shapiro.test(resid(alignment.lmer)) #p-value = 0.3076
```

```
##
##  Shapiro-Wilk normality test
##
## data:  resid(alignment.lmer)
## W = 0.98577, p-value = 0.3076
```

```
bartlett.test(resid(alignment.lmer), align$Groupsize) #p-value = 0.6501
```

```
##
##  Bartlett test of homogeneity of variances
##
## data:  resid(alignment.lmer) and align$Groupsize
## Bartlett's K-squared = 0.8614, df = 2, p-value = 0.6501
```

```
alignment.lmer2 = lmer(Alignment ~ Groupsize+Time + (1|Video), data=align, na.action =
"na.omit")
anova(alignment.lmer, alignment.lmer2) #similar AIC so keep the more complex model
```

```
## Data: align
## Models:
## alignment.lmer2: Alignment ~ Groupsize + Time + (1 | Video)
## alignment.lmer: Alignment ~ Groupsize * Time + (1 | Video)
##                  npar     AIC     BIC logLik deviance  Chisq Df Pr(>Chisq)
## alignment.lmer2    7 -106.93 -88.151 60.463  -120.93
## alignment.lmer    11 -107.08 -77.579 64.541  -129.08 8.1573  4    0.08598 .
## ---
## Signif. codes:  0 '***' 0.001 '**' 0.01 '*' 0.05 '.' 0.1 ' ' 1
```

```
summary(alignment.lmer)
```

```
## Linear mixed model fit by REML ['lmerMod']
## Formula: Alignment ~ Groupsize * Time + (1 | Video)
##    Data: align
##
## REML criterion at convergence: -87.3
##
## Scaled residuals:
##     Min      1Q  Median      3Q     Max
## -2.3561 -0.5491  0.1028  0.5077  1.7859
##
## Random effects:
##  Groups   Name        Variance Std.Dev.
##  Video    (Intercept) 0.01331  0.1154
##  Residual             0.01182  0.1087
## Number of obs: 108, groups:  Video, 36
##
## Fixed effects:
##                     Estimate Std. Error t value
## (Intercept)          0.93517    0.04575  20.439
## Groupsize8          -0.06975    0.06471  -1.078
## Groupsize16         -0.18333    0.06471  -2.833
## Time30              -0.23083    0.04438  -5.202
## Time100             -0.20592    0.04438  -4.640
## Groupsize8:Time30    0.06092    0.06276   0.971
## Groupsize16:Time30   0.15483    0.06276   2.467
## Groupsize8:Time100   0.09500    0.06276   1.514
## Groupsize16:Time100  0.13667    0.06276   2.178
##
## Correlation of Fixed Effects:
##             (Intr) Grpsz8 Grps16 Time30 Tim100 G8:T30 G16:T3 G8:T10
## Groupsize8  -0.707
## Groupsize16 -0.707  0.500
## Time30      -0.485  0.343  0.343
## Time100     -0.485  0.343  0.343  0.500
## Grpsz8:Tm30  0.343 -0.485 -0.242 -0.707 -0.354
## Grpsz16:T30  0.343 -0.242 -0.485 -0.707 -0.354  0.500
## Grpsz8:T100  0.343 -0.485 -0.242 -0.354 -0.707  0.500  0.250
## Grps16:T100  0.343 -0.242 -0.485 -0.354 -0.707  0.250  0.500  0.500
```

```
Anova(alignment.lmer, test="F")
```

```
## Analysis of Deviance Table (Type II Wald F tests with Kenward-Roger df)
##
## Response: Alignment
##                      F Df Df.res    Pr(>F)
## Groupsize       1.4402  2     33    0.2514
## Time           21.6978  2     66 5.727e-08 ***
## Groupsize:Time  1.9794  4     66    0.1079
## ---
## Signif. codes:  0 '***' 0.001 '**' 0.01 '*' 0.05 '.' 0.1 ' ' 1
```

```
r.squaredGLMM(alignment.lmer)
```

```
##            R2m       R2c
## [1,] 0.2193555 0.6328307
```

```
emmeans(alignment.lmer, pairwise~Time, adjust = c("Tukey"))
```

```
## $emmeans
##  Time emmean     SE   df lower.CL upper.CL
##  0     0.851 0.0264 63.4    0.798    0.904
##  30    0.692 0.0264 63.4    0.639    0.745
##  100   0.722 0.0264 63.4    0.669    0.775
##
## Results are averaged over the levels of: Groupsize
## Degrees-of-freedom method: kenward-roger
## Confidence level used: 0.95
##
## $contrasts
##  contrast         estimate     SE df t.ratio p.value
##  Time0 - Time30     0.1589 0.0256 66   6.203  <.0001
##  Time0 - Time100    0.1287 0.0256 66   5.023  <.0001
##  Time30 - Time100  -0.0302 0.0256 66  -1.180  0.4695
##
## Results are averaged over the levels of: Groupsize
## Degrees-of-freedom method: kenward-roger
## P value adjustment: tukey method for comparing a family of 3 estimates
```

```
emmeans(alignment.lmer, pairwise~Groupsize*Time, adjust = c("Tukey"))
```

```
## $emmeans
##  Groupsize Time emmean     SE   df lower.CL upper.CL
##  4         0     0.935 0.0458 63.4    0.844    1.027
##  8         0     0.865 0.0458 63.4    0.774    0.957
##  16        0     0.752 0.0458 63.4    0.660    0.843
##  4         30    0.704 0.0458 63.4    0.613    0.796
##  8         30    0.696 0.0458 63.4    0.604    0.787
##  16        30    0.676 0.0458 63.4    0.584    0.767
##  4         100   0.729 0.0458 63.4    0.638    0.821
##  8         100   0.754 0.0458 63.4    0.663    0.846
##  16        100   0.683 0.0458 63.4    0.591    0.774
##
## Degrees-of-freedom method: kenward-roger
## Confidence level used: 0.95
##
## $contrasts
##  contrast                                   estimate     SE   df t.ratio p.value
##  Groupsize4 Time0 - Groupsize8 Time0         0.06975 0.0647 63.4   1.078  0.9754
##  Groupsize4 Time0 - Groupsize16 Time0        0.18333 0.0647 63.4   2.833  0.1261
##  Groupsize4 Time0 - Groupsize4 Time30        0.23083 0.0444 66.0   5.202  0.0001
##  Groupsize4 Time0 - Groupsize8 Time30        0.23967 0.0647 63.4   3.704  0.0124
##  Groupsize4 Time0 - Groupsize16 Time30       0.25933 0.0647 63.4   4.008  0.0048
##  Groupsize4 Time0 - Groupsize4 Time100       0.20592 0.0444 66.0   4.640  0.0005
##  Groupsize4 Time0 - Groupsize8 Time100       0.18067 0.0647 63.4   2.792  0.1383
##  Groupsize4 Time0 - Groupsize16 Time100      0.25258 0.0647 63.4   3.903  0.0067
##  Groupsize8 Time0 - Groupsize16 Time0        0.11358 0.0647 63.4   1.755  0.7103
##  Groupsize8 Time0 - Groupsize4 Time30        0.16108 0.0647 63.4   2.489  0.2568
##  Groupsize8 Time0 - Groupsize8 Time30        0.16992 0.0444 66.0   3.829  0.0083
##  Groupsize8 Time0 - Groupsize16 Time30       0.18958 0.0647 63.4   2.930  0.1009
##  Groupsize8 Time0 - Groupsize4 Time100       0.13617 0.0647 63.4   2.104  0.4797
##  Groupsize8 Time0 - Groupsize8 Time100       0.11092 0.0444 66.0   2.499  0.2513
##  Groupsize8 Time0 - Groupsize16 Time100      0.18283 0.0647 63.4   2.826  0.1283
##  Groupsize16 Time0 - Groupsize4 Time30       0.04750 0.0647 63.4   0.734  0.9981
##  Groupsize16 Time0 - Groupsize8 Time30       0.05633 0.0647 63.4   0.871  0.9938
##  Groupsize16 Time0 - Groupsize16 Time30      0.07600 0.0444 66.0   1.713  0.7367
##  Groupsize16 Time0 - Groupsize4 Time100      0.02258 0.0647 63.4   0.349  1.0000
##  Groupsize16 Time0 - Groupsize8 Time100     -0.00267 0.0647 63.4  -0.041  1.0000
##  Groupsize16 Time0 - Groupsize16 Time100     0.06925 0.0444 66.0   1.560  0.8221
##  Groupsize4 Time30 - Groupsize8 Time30       0.00883 0.0647 63.4   0.137  1.0000
##  Groupsize4 Time30 - Groupsize16 Time30      0.02850 0.0647 63.4   0.440  1.0000
##  Groupsize4 Time30 - Groupsize4 Time100     -0.02492 0.0444 66.0  -0.561  0.9997
##  Groupsize4 Time30 - Groupsize8 Time100     -0.05017 0.0647 63.4  -0.775  0.9972
##  Groupsize4 Time30 - Groupsize16 Time100     0.02175 0.0647 63.4   0.336  1.0000
##  Groupsize8 Time30 - Groupsize16 Time30      0.01967 0.0647 63.4   0.304  1.0000
##  Groupsize8 Time30 - Groupsize4 Time100     -0.03375 0.0647 63.4  -0.522  0.9998
##  Groupsize8 Time30 - Groupsize8 Time100     -0.05900 0.0444 66.0  -1.330  0.9187
##  Groupsize8 Time30 - Groupsize16 Time100     0.01292 0.0647 63.4   0.200  1.0000
##  Groupsize16 Time30 - Groupsize4 Time100    -0.05342 0.0647 63.4  -0.826  0.9956
##  Groupsize16 Time30 - Groupsize8 Time100    -0.07867 0.0647 63.4  -1.216  0.9500
##  Groupsize16 Time30 - Groupsize16 Time100 -0.00675 0.0444 66.0  -0.152  1.0000
##  Groupsize4 Time100 - Groupsize8 Time100    -0.02525 0.0647 63.4  -0.390  1.0000
##  Groupsize4 Time100 - Groupsize16 Time100    0.04667 0.0647 63.4   0.721  0.9983
```

```
##  Groupsize8 Time100 - Groupsize16 Time100  0.07192 0.0647 63.4   1.111  0.9704
##
## Degrees-of-freedom method: kenward-roger
## P value adjustment: tukey method for comparing a family of 9 estimates
```

```
alignment.emm <- ref_grid(alignment.lmer, ~Groupsize*Time, type="response")
summary(alignment.emm)
```

```
##  Groupsize Time prediction     SE   df
##  4         0         0.935 0.0458 63.4
##  8         0         0.865 0.0458 63.4
##  16        0         0.752 0.0458 63.4
##  4         30        0.704 0.0458 63.4
##  8         30        0.696 0.0458 63.4
##  16        30        0.676 0.0458 63.4
##  4         100       0.729 0.0458 63.4
##  8         100       0.754 0.0458 63.4
##  16        100       0.683 0.0458 63.4
##
## Degrees-of-freedom method: kenward-roger
```

```
alignment.emm<-as.data.frame(alignment.emm)
align.plot = ggplot(alignment.emm, aes(x=Time, y=prediction, fill=Groupsize)) +
  geom_pointrange(aes(ymin = prediction - SE, ymax = prediction + SE, color = Groupsiz
e), na.rm = TRUE, shape=21, size=.7, data = alignment.emm) +
  geom_line(aes(group = Groupsize, colour = Groupsize), na.rm = TRUE, alignment.emm) +
  geom_jitter(aes(x=Time, y = Alignment, colour=Groupsize), shape = 21, size=1.2, width
=.25, alpha = 0.3, data = align) +
  xlab(bquote('Time post-stimulus (ms)')) + ylab(bquote('Alignment (r)')) +
  theme_bw() + theme(panel.grid.major = element_blank(), panel.grid.minor = element_blan
k()) +
  theme(axis.text = element_text(size = 12)) + theme(axis.title = element_text(size = 1
2)) +
  theme_classic(base_family='Arial', base_size = 28) +
  theme(legend.title=element_blank()) +
  theme(legend.position = "top")

align.plot
```

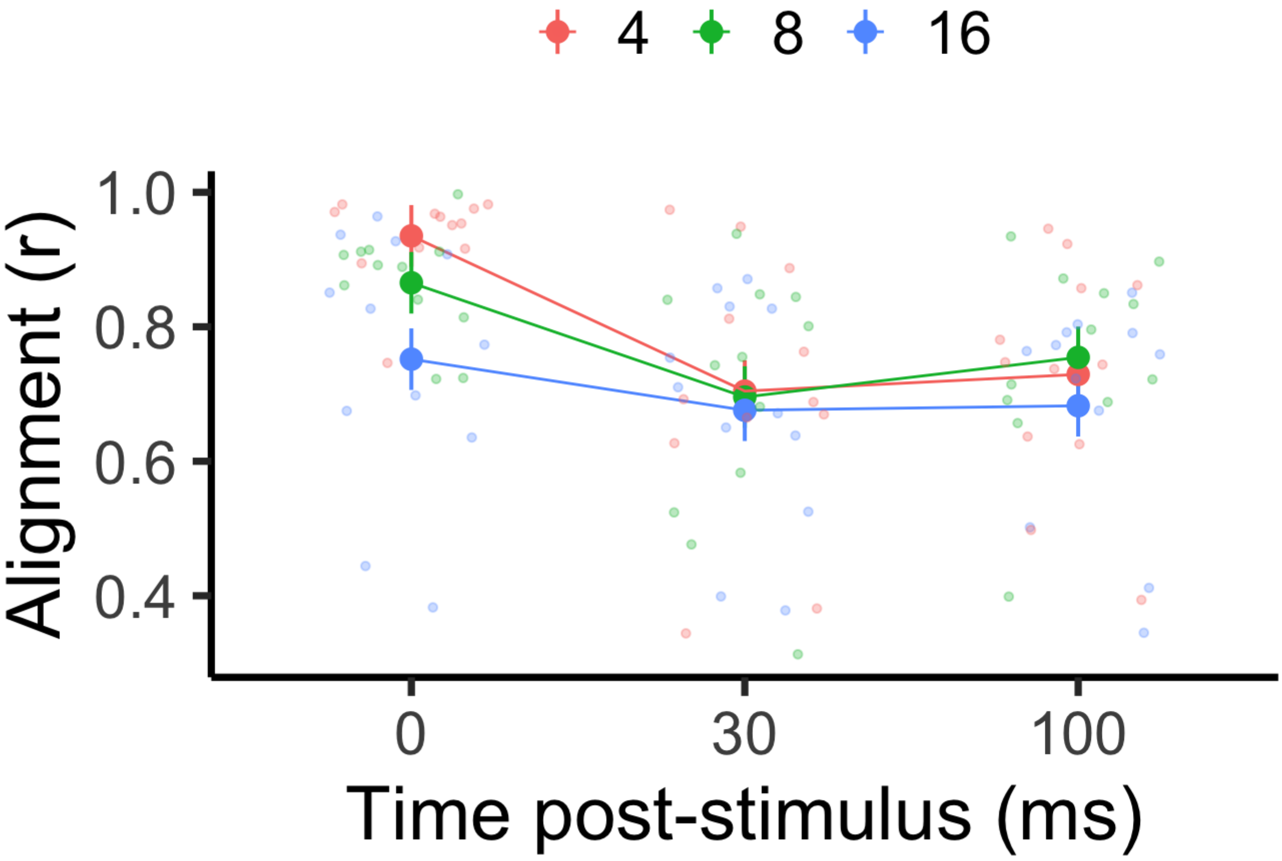
4
8
16
Alignment (r)
1.0
0.8
0.6
0.4
0
30
100
Time post-stimulus (ms)
